# Genome-scale chromatin interaction dynamic measurements for key components of the RNA Pol II general transcription machinery

**DOI:** 10.1101/2023.07.25.550532

**Authors:** Kristyna Kupkova, Savera J. Shetty, Elizabeth A. Hoffman, Stefan Bekiranov, David T. Auble

## Abstract

**Background:** A great deal of work has revealed in structural detail the components of the machinery responsible for mRNA gene transcription initiation. These include the general transcription factors (GTFs), which assemble at promoters along with RNA Polymerase II (Pol II) to form a preinitiation complex (PIC) aided by the activities of cofactors and site-specific transcription factors (TFs). However, less well understood are the *in vivo* PIC assembly pathways and their kinetics, an understanding of which is vital for determining on a mechanistic level how rates of *in vivo* RNA synthesis are established and how cofactors and TFs impact them.

**Results:** We used competition ChIP to obtain genome-scale estimates of the residence times for five GTFs: TBP, TFIIA, TFIIB, TFIIE and TFIIF in budding yeast. While many GTF-chromatin interactions were short-lived (< 1 min), there were numerous interactions with residence times in the several minutes range. Sets of genes with a shared function also shared similar patterns of GTF kinetic behavior. TFIIE, a GTF that enters the PIC late in the assembly process, had residence times correlated with RNA synthesis rates.

**Conclusions:** The datasets and results reported here provide kinetic information for most of the Pol II-driven genes in this organism and therefore offer a rich resource for exploring the mechanistic relationships between PIC assembly, gene regulation, and transcription. The relationships between gene function and GTF dynamics suggest that shared sets of TFs tune PIC assembly kinetics to ensure appropriate levels of expression.

## Background

Transcription is a highly complex biochemical process whose exquisite regulation is of fundamental importance in determining cell function and fate. A tremendous amount of information is available on the structure, biochemical functions, and relationships of various transcription factors (TFs), co-factors, and subunits of the general transcription machinery (1–5). This includes structures of the transcription preinitiation complex (PIC), which assembles at promoters and consists of the general transcription factors (GTFs) TFIIA, TFIIB, TFIID, TFIIE, TFIIF, and TFIIH, as well as RNA polymerase II (Pol II) (1–3,6–9). In addition, genome-wide analyses have provided global snapshots of many factors along the eukaryotic DNA template (10–13). These combined studies have led to a conceptual framework in which PICs are assembled stepwise at promoters. This process begins with nucleation by TFIID, a multisubunit complex that contains the DNA-binding subunit TATA-binding protein (TBP) (14,15), and can be further facilitated by binding of TFs and coactivators that physically contact GTFs (16). *In vitro*, following the binding of TBP/TFIID to a TATA-containing promoter, TFIIA and TFIIB can then associate with the complex, followed by Pol II in association with TFIIF, and then TFIIE (17). This multi-subunit complex provides the substrate for recruitment of TFIIH (18), whose activities are required *in vivo* but may be dispensable *in vitro* using naked DNA substrates (19). A key factor contributing to PIC assembly *in vivo* is Mediator, which physically contacts multiple GTFs and modulates the activities of TFIIH (19–21). Live-cell imaging has documented the dynamic behavior of these factors and is generally consistent with such an assembly pathway, albeit occurring via highly dynamic and short-lived complexes (22). Importantly, the understanding of PIC assembly has emerged mainly from studies that have focused on the analysis of stable complexes formed *in vitro* or identified *in vivo*, lacking information about the locus-specific dynamics of the process. Furthermore, some evidence suggests that the canonical *in vitro* assembly pathway may not apply to PICs at all promoters *in vivo* (23–25). In addition to unexplored assembly pathway complexity, it has become apparent that *in vivo* transcription is a highly dynamic and stochastic process, with RNA synthesis often occurring from individual genes in bursts, and with variability occurring among genetically identical cells (26,27). Most models of RNA expression based on these types of observations do not posit particular features of protein-DNA complex behavior as the explanation, and relatively few genes have been analyzed in depth (28–32). Indeed, live cell imaging approaches have revealed that while TFs in general display very dynamic interactions with chromatin, the functional consequences of their interaction kinetics are only beginning to be explored on a mechanistic level (22,33,34).

The premise of this study is that PIC assembly dynamics are variable across the genome and that identification of kinetic pathways in PIC assembly will shed light on mechanisms of regulation that operate at the level of transcription initiation. To better understand PIC assembly *in vivo*, we have used an approach called competition chromatin immunoprecipitation (competition ChIP, ref.(35)) to measure the site-specific, genome-scale chromatin binding dynamics of five GTFs (TBP, TFIIA, TFIIB, TFIIE, and TFIIF) in the budding yeast *S. cerevisiae*. In addition, we compared promoter binding dynamics of these factors with RNA synthesis rates to determine how chromatin binding of key PIC components relates to the production of RNA. To our knowledge, this represents the first comprehensive analysis of PIC dynamics, provides a global picture of PIC assembly, and highlights promoter-specific variation.

## Results

Competition ChIP (CC) is an approach in which cells harbor two isoforms of a transcription factor of interest with distinguishable epitope tags (Fig. 1a). We engineered diploid yeast cells to constitutively express one isoform with a Myc tag under control of the endogenous promoter and with the second isoform tagged with HA and under inducible *GAL* promoter control. In the CC experiments, cells were shifted to galactose at time zero to induce expression of the HA-tagged competitor isoform, followed by cell culture sample collection at various time points (Fig. 1b). We then measured the relative occupancies of the Myc- and HA-tagged species genome-wide at each time point (Fig. 1c) and used the relative occupancies as input to a model that describes the competition for chromatin binding to each site, yielding the site-specific residence time (Fig. 1d). The principle of the assay is outlined in Fig. 1e,f, which illustrate how the occupancy ratios of the two isoforms of a particular factor would change if the factor has a short or long residence time at a particular site. Notably, TFIIA, TFIIE, and TFIIF are biochemically composed of more than one subunit, and thus, for these factors we epitope tagged one subunit and placed one copy of each subunit under *GAL* control in order to induce balanced expression when cells were grown in galactose (see Methods).

**Fig. 1.**
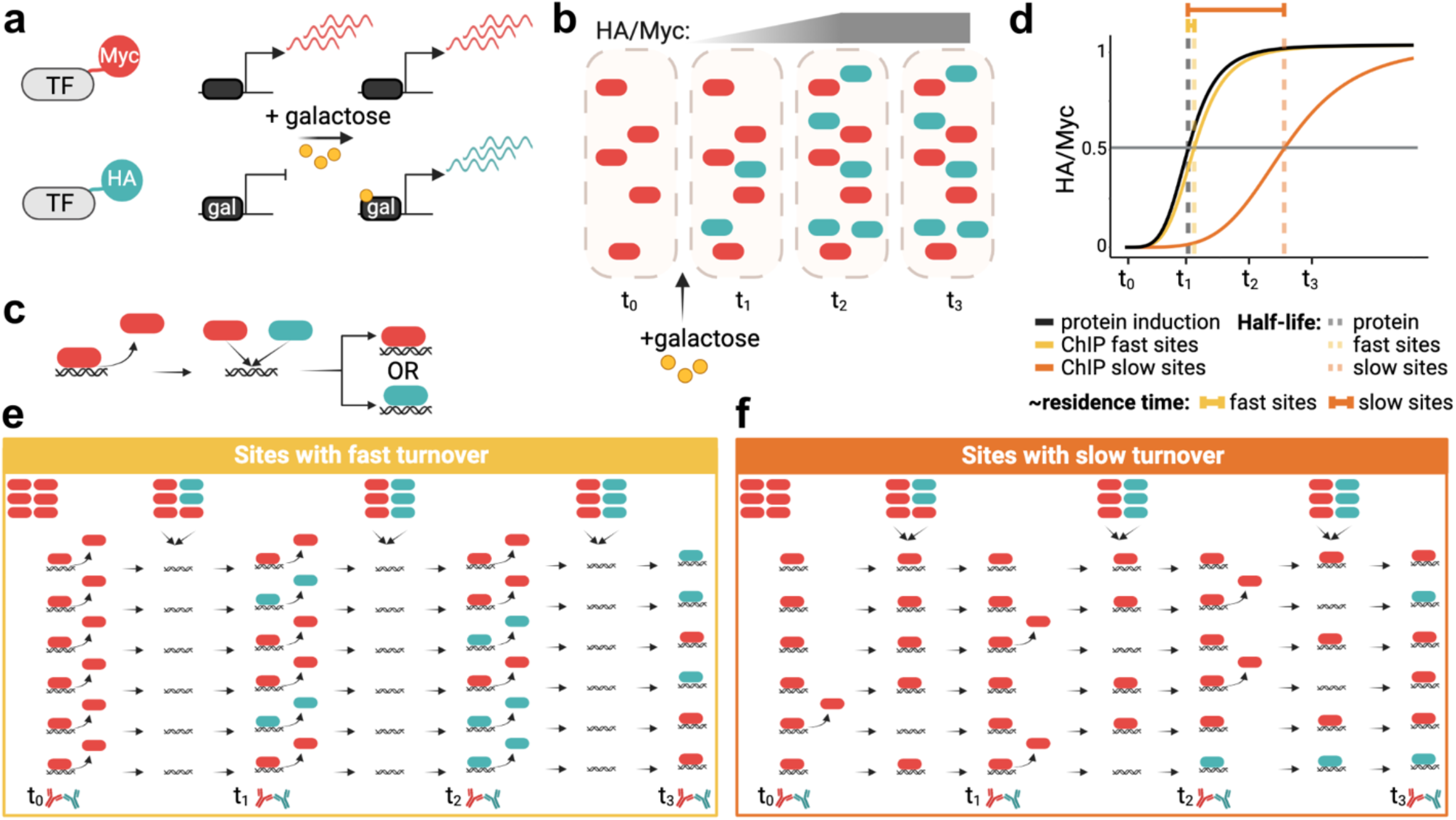
Competition ChIP overview. (a) A Myc-tagged isoform of a TF is expressed constitutively using the endogenous promoter, while an HA-isoform is expressed under control of a galactose-inducible promoter. (b) Illustration showing protein induction upon adding galactose. The HA/Myc ratio increases over time until it reaches saturation. (c) Once a given TF unbinds DNA, the two isoforms compete for binding to the available site. (d) Simplified illustration of residence time estimation based on the lag of the normalized HA/Myc ChIP signal ratio relative to the competitor protein induction curve, as further illustrated in (e) for sites with fast turnover and (f) for sites with slow turnover. In (e,f) the icons in the top row indicate relative levels of constitutive (red) and competitor (green) isoforms.

For each factor, we first measured the levels of both isoforms by Western blotting (Fig. 2a-c; Additional file 1: Fig. S1, Additional file 2: Table S1). The time-dependent accumulation of competitor isoforms could be fit to the Hill equation with induction half-times of ∼43 min and Hill coefficients of ∼4.5 on average (Fig. 2b,c). We estimated residence times by fitting the normalized time-dependent turnover ratios to a turnover model (36) (see Methods), and compared the fits to the HA-tagged competitor’s synthesis rate. In this way, we were able to assign residence times for binding interactions with significantly longer (> 1 min) rates of turnover compared to the rate of competitor synthesis, and for reliable fits that were not significantly different from the rate of competitor induction, we were able to classify the chromatin binding residence times as < 1 min (see Methods). Overall, we were able to estimate residence times for each GTF binding to ∼3000 or more promoters (Fig. 2d; Additional file 3: Table S2). This represents roughly half of the Pol II promoters in the *S. cerevisiae* genome. Representative fits are shown in Fig. 2e; Additional file 1: Fig. S2. Note that the HA/Myc ratios at sites with rapid turnover closely mimic the time course of competitor induction, whereas more long-lived complexes have turnover ratios that are notably displaced to the right of the competitor induction curves. The distributions of turnover times are shown in Fig. 2f. We identified different numbers of sites for each TF for which we were able to assign residence times; this is indicative of differences in the number of sites for which we were able to obtain reliable fits of the kinetic data, as well as likely differences in the efficiency of formaldehyde capture of short-lived complexes. It is notable that the majority of TBP, TFIIA, TFIIB, and TFIIF chromatin interactions were short-lived (i.e. < 1-2 min) whereas the majority of TFIIE complexes displayed residence times in the several minutes range. It was also notable that TFIIF residence times were bimodal, with most estimates being short-lived (∼2 min or less) and the rest in the 5-10 min range (Fig. 2f; discussed below).

**Fig. 2.**
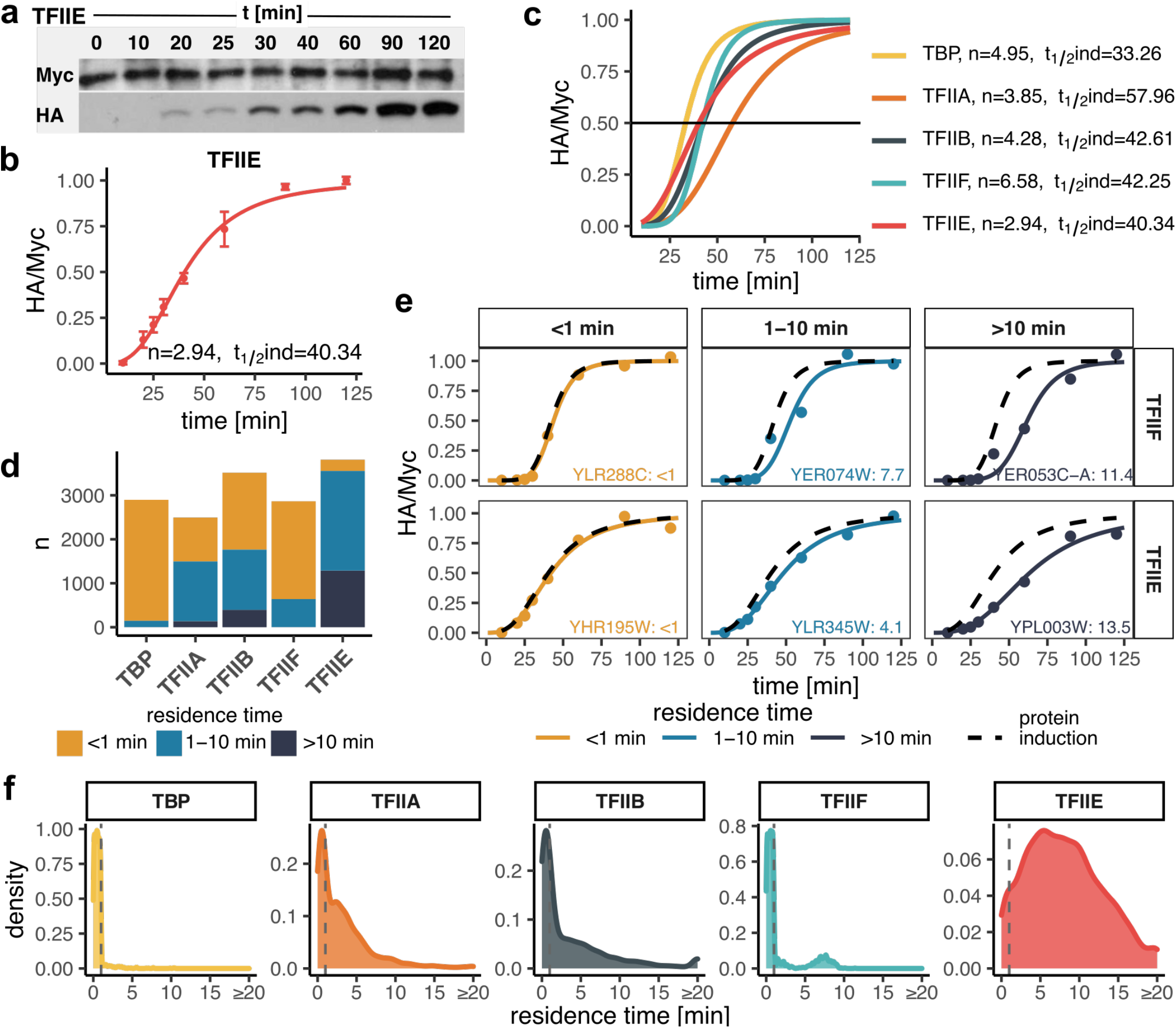
GTF residence times. (a) TFIIE Western blots showing the isoform levels of the TFIIE subunit over the indicated time course. Galactose was added at t = 0 min. (b) Quantified Western blots from (a). Shown are normalized HA/Myc ratios with error bars representing standard deviation (n = 3). The induction curve was fitted with a Hill coefficient (n) and induction half-time (t_½_ind) as indicated. (c) Induction curves as in (b) for all targeted TFs with fit parameters indicated on the right. (d) Bar plot showing the number of sites (y-axis) categorized based on estimated residence time for each TF (x-axis). (e) Examples of sites with fast (<1 min), moderate (1-10 min) and slow (>10 min) turnover for TFIIF and TFIIE. Black dashed curves represent protein induction curves from (c), in color are shown the normalized HA/Myc ChIP signals (mapped reads) along with the fitted model. Gene target names along with the estimated residence times are included. (f) Distribution of estimated residence times for all GTFs. Values for reliably fast sites (<1 min) were randomly generated for plotting purposes and are separated by dashed lines.

To determine the relationship between GTF promoter residence time and the rate of RNA synthesis from the corresponding genes, we measured newly synthesized RNA under these same conditions (Additional file 1: Fig. S3a; Additional file 4: Table S3). Replicate samples (n=2) were acquired at 20 and 60 minutes post galactose induction. There was excellent agreement between the replicates and between the two time points (Fig. 3a; Additional file 1: Fig. S3b-d). Dynamic transcriptome analysis (DTA, ref. (37)) was applied to estimate RNA synthesis rates (Additional file 1: Fig. S3e), which were in reasonable concordance with earlier data from cells grown in galactose (Additional file 1: Fig. S3f, ref. (38)). We divided the mRNA synthesis rates into quartiles and compared them to GTF residence times (Fig. 3b,c). Residence times for TFIIA and TFIIB were on average modestly shorter for highly expressed genes compared to genes with lower expression levels, which may suggest a kinetic bottleneck in PIC assembly for poorly expressed genes that occurs after the binding of these two factors (see Discussion). Strikingly, the average TFIIE residence time increased with gene expression level across these four groups of genes (Fig. 3c), suggesting that the TFIIE residence time is an indicator of gene expression level. To relate residence time to RNA synthesis more directly, we calculated the ratio of mRNA molecules made per GTF binding event, which we previously defined as transcription efficiency (TE, ref. (36)). TE was on average < 1 mRNA synthesized per binding event for TBP, TFIIA, TFIIB, and TFIIF (Fig. 3d), suggesting that binding events by these factors do not efficiently give rise to the synthesis of mRNA. In addition, the TE values increased gradually and progressively for these factors with TBP having the lowest TEs and TFIIE the highest, in line with the *in vitro* assembly pathway in which TBP binds to promoters first, followed by TFIIA and TFIIB, which provide a platform for binding of TFIIF in association with Pol II (17). Notably, the median TE for TFIIE was close to one, suggesting that binding of TFIIE to promoters was associated with the production of one mRNA molecule on average. The results suggest that PIC formation is an increasingly efficient process along a pathway from TBP to TFIIE, and that the assembly of a TFIIE-containing PIC is associated with the production of a single molecule of mRNA. Using all of the GTF residence time data for Principal Component Analysis (PCA) revealed a correlation between GTF binding dynamics and RNA synthesis along the first principal component, PC1 (Fig. 3e; Additional file 1: Fig. S4), which was driven mainly by the positive correlations between TFIIE/TFIIF and RNA synthesis rate (Fig. 3f,g). This conclusion was further supported by linear modeling of the GTF residence time contributions to transcription rates (Additional file 1: Fig. S5a,b).

**Fig. 3.**
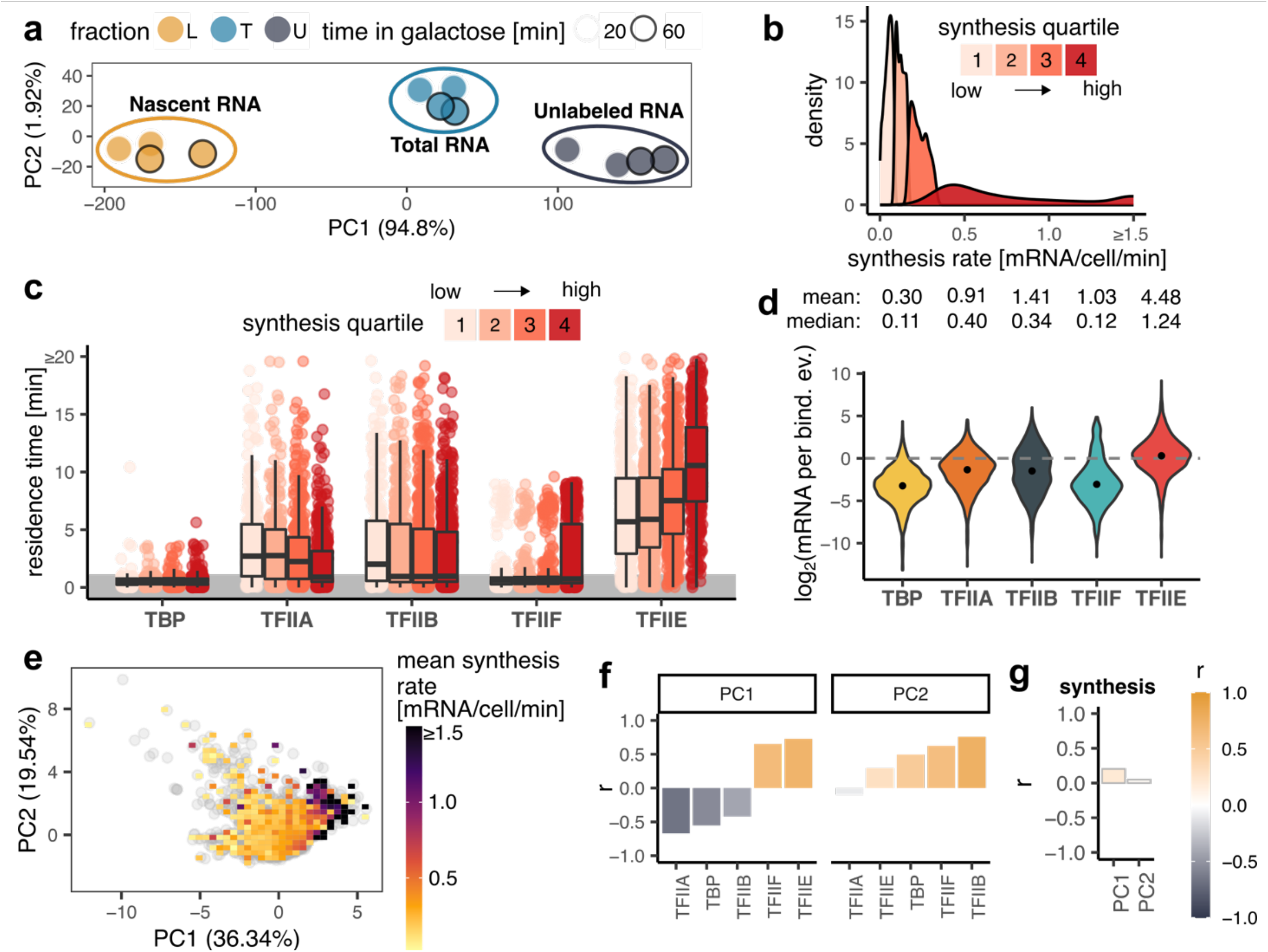
Relationship between residence GTF residence times and synthesis rates. (a) PCA plot showing low-dimensional representation of dynamic transcriptome analysis (DTA) samples without negative control. (b) Distribution of mRNA synthesis rate values separated into synthesis quartiles. (c) Box plots showing residence time distributions (yaxis) for all GTFs (x-axis) within the indicated synthesis quartile. Values for reliably fast sites (<1 min) were randomly generated for plotting purposes and are highlighted by the grey area. The middle line represents the median, the lower and upper edges of the boxes represent the first and third quartiles, and the whiskers represent 1.5 * interquartile range. (d) Violin plots showing the distributions of log_2_transformed transcription efficiency (TE, y-axis) for each GTF (x-axis). TE indicates the number of mRNA molecules synthesized during one binding event. The points show the medians of the log_2_ transformed TE values. Mean and median TE values are shown above the plots. (e) PCA plot showing low-dimensional representation of gene targets based on GTF residence times. Each grey point is a gene, color map shows the mean synthesis rate of genes under a given area. (f) Pearson’s correlation coefficients (y-axis) between the indicated PCs (panel title) and GTF (x-axis) residence times. (g) Pearson’s correlation coefficients (y-axis) between PCs (x-axis) and synthesis rates. In the PCA plots, the percentage within the axis labels indicates the percentage of variance explained by a given PC.

We next looked for pairwise relationships between the chromatin binding residence times of each GTF, and highlighted each gene by transcription rate (Fig. 4). TBP was less informative as most TBP binding events measured were short-lived and not well correlated with transcription rate (Additional file 1: Fig. S5c). In fact, the residence times of TFIIA, TFIIB, and TFIIF were not correlated with transcription rate either. This was in contrast to the positive correlation that was observed between TFIIE residence time and transcription rate (Additional file 1: Fig. S5). Interestingly, we observed a cluster of highly expressed genes whose promoters had long-lived TFIIE along with long-lived TFIIF.

**Fig. 4.**
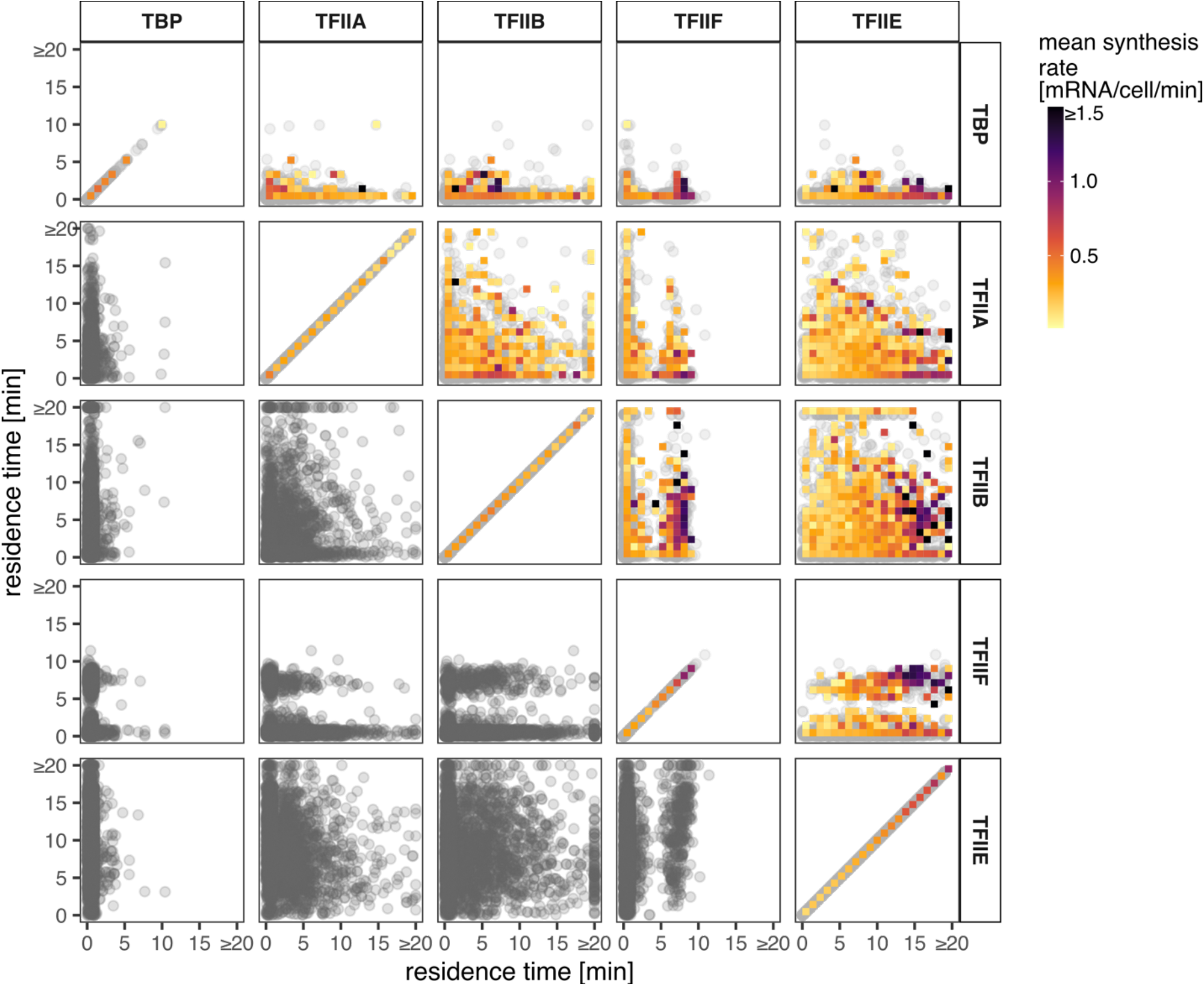
Relationships among GTF residence times and to mRNA synthesis rates. Each panel shows a comparison of residence times of pairs of GTFs as indicated in the panel titles. Each point is a shared gene target. The color map shows the mean synthesis rates of the genes under the given area.

Next, we clustered all the genes for which we obtained residence time measurements for four factors (TFIIA, TFIIB, TFIIE, and TFIIF (n = 1417)). We omitted TBP from this analysis due to the reduced number of sites with reliable estimated residence times > 1 min. We identified ten clusters spanning the full range of transcription rates (Fig. 5a,b; Additional file 5: Table S4).

**Fig. 5.**
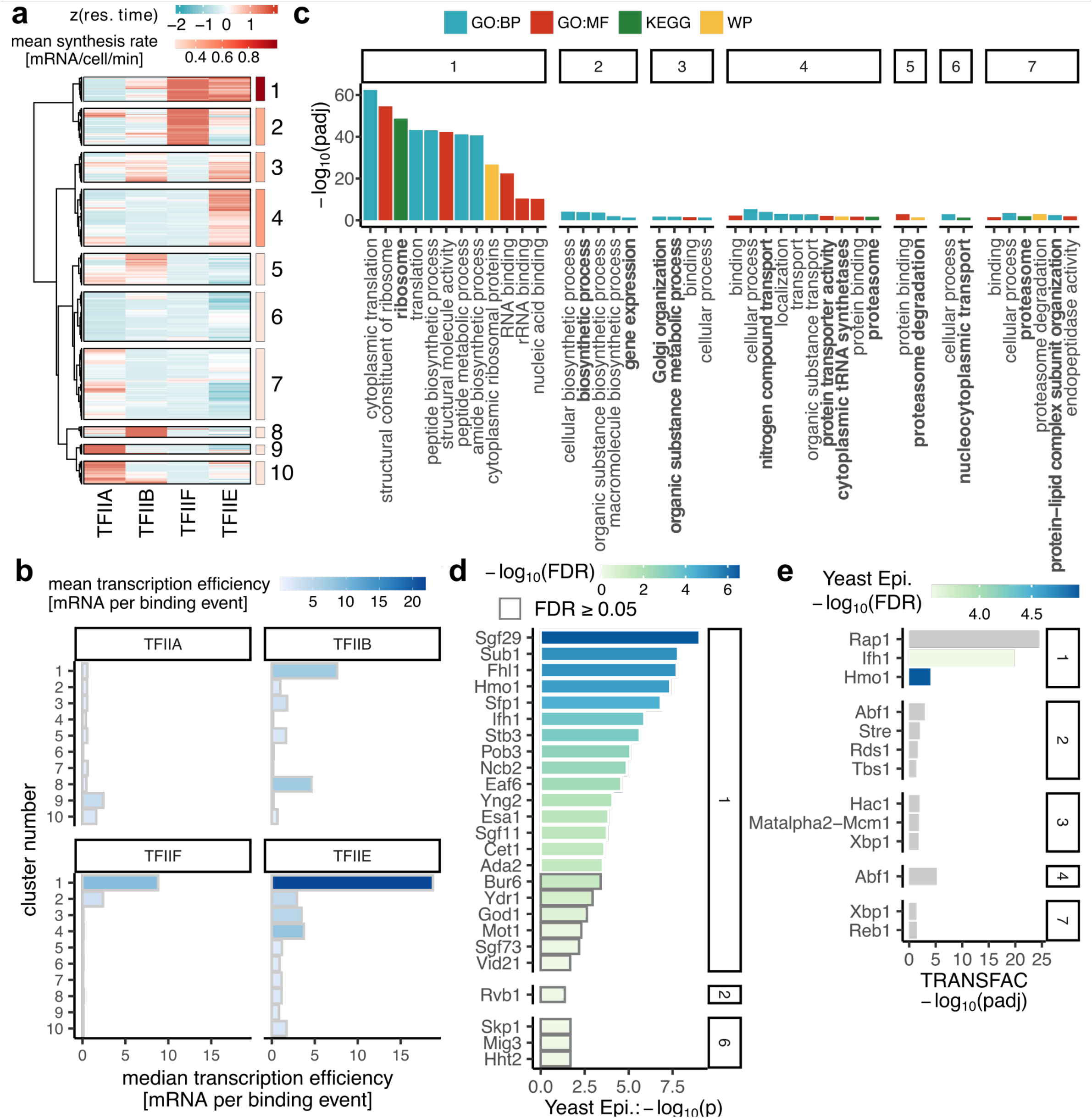
Gene classes based on GTF residence times combinations. (a) Heatmap showing z-score normalized residence times of the indicated GTFs (columns) across gene targets (rows) with the available residence time estimations from all four GTFs (n = 1417). Colored panels on the right side show the mean synthesis rates of genes belonging to the ten clusters. (b) Bar plots showing median TE (x-axis) within clusters (y-axis) from (a) and color-coded based on mean TE. (c-e) Functional annotation of genes from clusters in (a). Cluster number is indicated in the panel titles. (c) Pathway enrichment. Padj < 0.05. (d) Yeast Epigenome database DBF enrichment excluding subunits of GTFs and Pol II. P < 0.05. (e) TRANSFAC enrichment. Padj < 0.05. Colored bars were identified as significantly enriched (FDR < 0.05) in the Yeast Epigenome database.

Consistent with the results presented above, the most highly expressed genes had longer-lived TFIIE and/or TFIIF, whereas poorly expressed genes had promoters with longer-lived TFIIA. Longer residence times of TFIIB were associated with genes in several clusters, including in particular genes that were poorly expressed (cluster 8; Fig. 5a,b). Notably, the relatively long residence times of both TFIIE and TFIIF at cluster 1 promoters were associated with the production of multiple mRNAs, suggesting the formation of stable (sub)complexes that promote transcriptional bursting. In support of the biological significance of the observed residence time differences, genes within clusters 1-7 were functionally related (Fig. 5c). Cluster 1 genes include ribosomal protein genes and genes involved in RNA binding and translation. Additionally, cluster 2 genes are involved in biosynthetic processes; cluster 3 genes include those involved in Golgi organization; cluster 4 genes are involved in localization, transport, and the proteasome; cluster 5 genes are involved in proteasome degradation; cluster 6 genes have roles in nucleocytoplasmic transport; and cluster 7 genes are involved in proteasome and protein-lipid complex organization. The longer GTF residence times (as well as higher gene expression rates) at ribosomal protein genes in cluster 1 compared to the GTF residence times at other genes are statistically highly significant (Additional file 1: Fig. S6a-c). Moreover, expression of genes in most of these clusters is controlled by particular TFs (or sets of TFs; Fig. 5d,e, Additional file 1: Fig. S7), suggesting a mechanistic relationship between particular TFs and PIC assembly dynamics. Modest but significant increases in TFIIA and TFIIE residence times were observed at promoters with strong TATA elements versus those without such an element (39); these changes were consistent with a significant increase in RNA synthesis rate driven by TATA-containing promoters versus those without strong TATA elements (Additional file 1: Fig. S8a-c).

An unexpected observation was the above mentioned bimodal distribution of TFIIF residence times (Fig. 2f, Fig. 4). We observed functional enrichment of the genes in each of these two classes, with promoters in both classes associated with different subsets of genes involved in translation/ribosome. Genes with short-lived TFIIF were further associated with other biosynthetic processes (Additional file 1: Fig. S9a,b). Consistent with this, the genes in the long-lived and short-lived TFIIF classes were associated with particular enriched TFs, some of which were shared (Additional file 1: Fig. S9c-f). Among the TFs associated with genes in the long- and short-lived TFIIF classes, Rap1 was of particular interest as competition ChIP data were available for Rap1 from a prior study (40). Although Rap1 residence times were not correlated with residence times for TBP or TFIIB, there was a moderate correlation between Rap1 residences times and the residence times for TFIIA and TFIIE (Pearson’s correlation coefficients ∼ 0.37 and 0.3, respectively), and Rap1 residence times were significantly longer at genes with long-lived TFIIF compared to genes with short-lived TFIIF (Additional file 1: Fig. S10a,b).

## Discussion

The computational approach employed here for extraction of kinetic parameters from CC data is well supported by comparison with previous work. The TBP residence times obtained by analysis of CC data in this study were correlated with the residence times obtained from an older study using microarray data (Additional file 1: Fig. S10c; (41)) and are also broadly consistent with kinetic results for TBP in human cells (42). This includes the rank ordering in which tRNA genes had much longer residence times than mRNA genes. Previously, we used a formaldehyde crosslinking kinetic approach, called CLK, to measure chromatin binding dynamics (43). While the CLK method is technically challenging as well as locus-specific (44), we observed a rough agreement between the kinetic parameters obtained by the two methods for the handful of loci for which complementary measurements are available (Additional file 1: Fig. S10d).

Live cell imaging has revealed that the majority of TF-chromatin interactions studied are short-lived, with residence times on the order of seconds (22,45– 51). This includes TFIIB (22,52,53), for which CC results are reported here. The observation of highly dynamic binding by TFs has led to the view that such dynamics enable temporally responsive regulation of gene expression, and that TF residence times are associated with the duration of bursts in which more than one RNA molecule is synthesized during the TF period of occupancy on the promoter (30,54–56). Consistent with the observation of frequent short-lived chromatin interactions for TFs, we observed that the majority of the interactions between TBP, TFIIA, TFIIB, or TFIIF and chromatin had residence times of less than one minute (Fig. 2f). It was not possible to reliably estimate the residence times of these short-lived interactions using CC, but they must last long enough to be captured by crosslinking. It is likely that other very short-lived interactions were not detectable by our method because of their inability to be crosslinked. Conversely, it is possible that long-lived chromatin interactions such as those we report here would be difficult to detect with live cell imaging particularly if they occur infrequently, although evidence is emerging for TF-chromatin binding residence times on the minutes time scale using live cell imaging (57).

The biological significance of the residence times reported here is supported by the functional enrichment of genes in each of the clusters (Fig. 5c). This argues strongly that GTF residence time dynamics are tuned to facilitate expression levels that ensure that cells function and respond in physiologically appropriate ways. Since these gene sets are controlled by specific sets of TFs (Fig. 5d,e), it is reasonable to suggest that GTF dynamics are influenced in predictable ways by the TFs that control expression of the associated genes. It is understood that TFs exert context-specific effects on gene expression, and such effects have been generally described in terms of effects mediated by co-regulatory interactions with other TFs as well as epigenetic control, including DNA methylation (58–60). In future work, it could be interesting to explore how GTF residence times are impacted by manipulation of such regulators. We suggest that RNA output resulting from the interplay of these variables is at least partly a consequence of the capacity to catalyze the formation of functional PICs by overcoming kinetic bottlenecks in PIC assembly that are also related to the underlying DNA sequence and chromatin environment.

A striking observation from the results of this study is that the residence time of TFIIE is correlated with the mRNA synthesis rate, and the ratio of mRNA molecules produced to the TFIIE residence time suggests that one TFIIE binding event is associated with the production of one mRNA molecule (Fig. 3d). This is in contrast to the other GTFs for which one binding event was associated with less than one mRNA molecule produced. We were not able to measure Pol II directly using CC because we do not have a system for inducing the expression of all of the Pol II subunits to generate a competitor isoform of Pol II. However, TFIIF can serve as a proxy for Pol II itself as biochemical and structural data support a model in which TFIIF enters the PIC in association with Pol II (3,61–64). The combined results suggest that the formation of a PIC is an inefficient process in vivo, with most interactions of GTFs leading to subcomplexes that decay rather than leading to formation of a PIC capable of producing mRNA. This general view of transcription initiation inefficiency is consistent with live cell imaging data obtained by analysis of a gene array in a mouse cell line (65). Moreover, this pattern is broadly consistent with a PIC assembly pathway derived from in vitro studies in which TBP/TFIID initially interacts with DNA directly, followed by the binding of TFIIA and TFIIB, which provide a platform for the binding of Pol II and TFIIF, and subsequently TFIIE (Fig. 6; (24)). We infer the existence of stable TFIIB complexes on the basis of slow turnover at a relatively small number of genes; it appears that most TFIIB-containing complexes are unstable and that assembly of TFIIB in the PIC requires Pol II (22). Despite the dispensability of some GTFs in vitro under certain conditions, our results are also consistent with depletion experiments showing that all of the GTFs are required for all Pol II-mediated transcription in vivo, and that stable, partially assembled PICs are not detectable (66). Of note, however, we did observe a small number of relatively long-lived complexes containing TFIIA or TFIIB (Fig. 2d,f; Table S1). Such long-lived complexes could be consistent with the formation of a subcomplex of GTFs that is durably bound to promoters and promotes reinitiation (67). The formation of long-lived scaffolds of GTFs at some promoters is also suggested by the residence times of TFIIE and TFIIF at Cluster 1 genes, which were associated with the production of multiple mRNAs (Fig. 5b). Lastly, our analysis includes the minimal set of GTFs required for in vitro transcription using a naked DNA template (68–70). In future work and using methods suitable for analysis of multi-subunit complexes, it will be interesting to investigate the dynamics of TFIIH (71,72), Mediator and Pol II itself (73,74). Other important questions that could be addressed by performing kinetic measurements in suitably perturbed cells include probing the roles of promoter chromatin structure, and particularly the function of the first nucleosome (66). Taken together, we feel that the results presented here provide a foundation for future work to understand how TFs, co-factors, and the native chromatin environment contribute mechanistically to the establishment of the rates of transcription initiation observed *in vivo*.

**Fig. 6.**
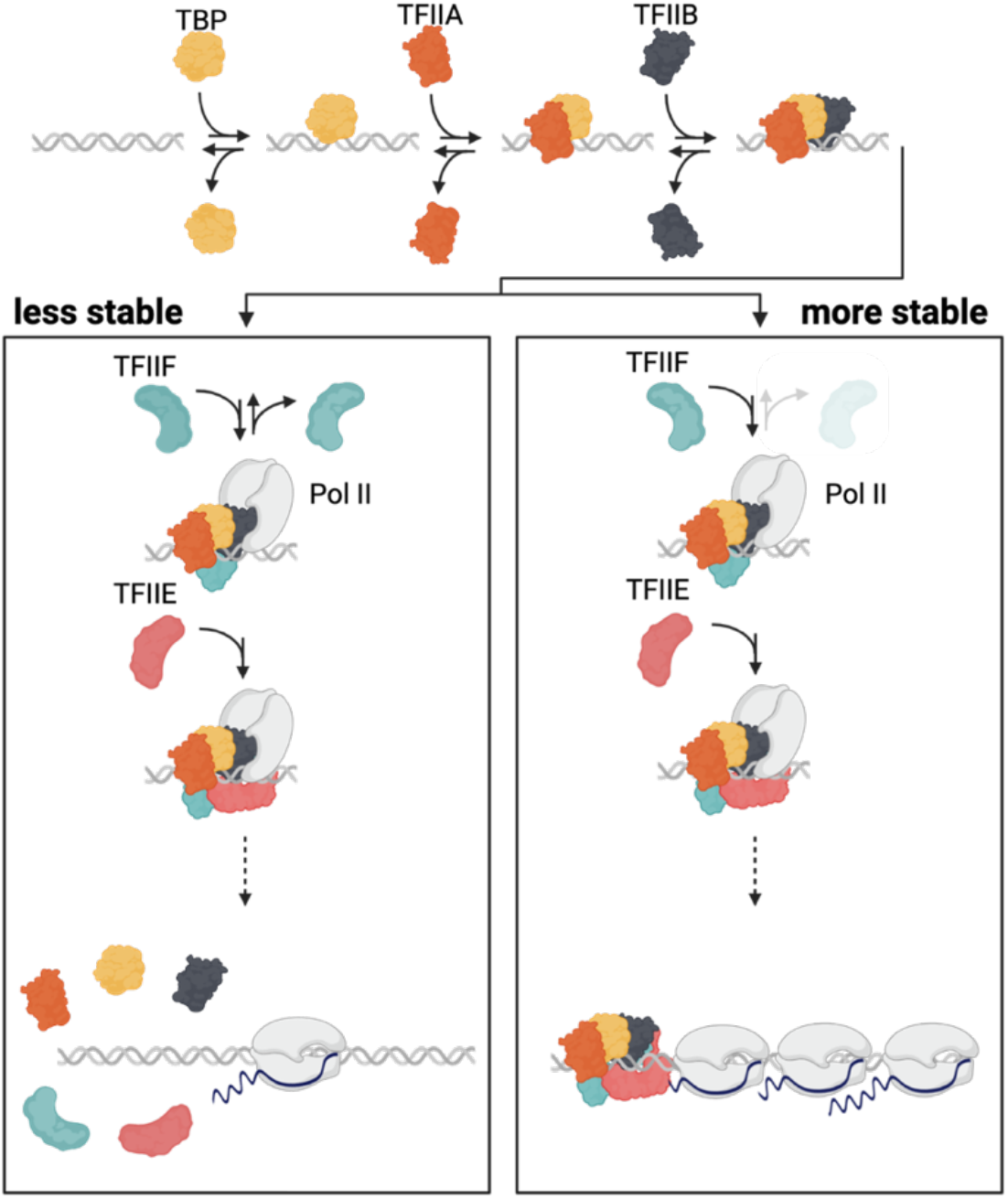
Model. The results presented here suggest that for the majority of genes, PICs are unstable until TFIIE binding which leads to functional PIC assembly, the initiation of RNA synthesis and release of Pol II and PIC disassembly. At a relatively small subset of genes e.g., genes coding for ribosomal subunits, relatively stable PICs are formed upon TFIIF binding (note lighter color for disassociation) and further stabilized by TFIIE binding, followed by the initiation of RNA synthesis. Upon Pol II release, stable PICs may be disassembled or at certain promoters may be stable and lead to transcriptional bursting. The formation of more stable PICs is likely associated with promoter-specific features and cofactors. The figure is meant to be illustrative and does not represent accurate sizes or molecular shapes of the factors of interest.

## Conclusions

The results reported here provide a wealth of kinetic information describing the chromatin binding dynamics of five key GTFs at the majority of promoters in budding yeast. In general agreement with live cell imaging results, we find that many interactions are too short-lived to be measured by CC. However, there are many interactions with residence times in the several minutes range, and importantly, promoters with shared GTF kinetics are functionally related. This supports a model in which the rates of RNA synthesis *in vivo* are influenced or perhaps controlled by rates of PIC assembly, which themselves result from the combination of promoter sequence, chromatin environment and the TFs and cofactors that impact them. Overall, the kinetic behavior is consistent with the stepwise PIC assembly pathway established using purified components *in vitro* and in which the RNA synthesis rate is closely correlated with the residence time of TFIIE. These results suggest that at most promoters, relatively unstable GTF subcomplexes give rise to more stable fully assembled PICs and that the initiation of RNA synthesis is accompanied by PIC dissolution. At certain promoters, GTF binding events are associated with the production of multiple mRNAs, suggesting the formation of stable PIC subcomplexes that facilitate transcription reinitiation.

## Methods

### Yeast strains

The parental diploid strain W303 (75) was used to generate all of the competition ChIP strains. For each GTF, one allele was N-terminally tagged with 3xHA and placed under the control of an inducible *GAL1* promoter. The other allele was N-terminally tagged with 9xMyc and remained under the control of the endogenous promoter (76).

For construction of the *GAL1*-induced alleles, the plasmid pFA6-His3MX6-PGAL1-3HA (RRID:Addgene_41610, ref. (76)) was used to obtain the His3MX6-PGAL1-3HA cassette by PCR amplification (see Additional file 6: Table S5 for primers) and was integrated into the genome using standard yeast molecular biology techniques. For the GTFs TFIIA, TFIIE, and TFIIF, which consist of two subunits, one copy of each subunit was placed under *GAL1* control to ensure balanced expression of the competitor isoform. Following integration of the *HIS3-GAL1*-3HA cassette at one gene subunit, the strain was transformed with the *TRP1-GAL1* cassette from pFA6-TRP1-PGAL1 (RRID:Addgene_41606, ref. (76)), placing the second subunit under *GAL1* control but without an epitope tag. The 9xMyc tag was integrated into the genome of another isolate of W303 using the integration and Cre-recombinase knockout method and reagents developed by Gauss et al (77). The 9xMyc tag and loxP-flanked KanMX6 marker were PCR amplified from pOM20 and integrated into the yeast genome using standard methods as above. The KanMX6 marker was then then deleted using the *GAL-* inducible Cre recombinase carried on the plasmid pSH47 (78). The Myc-tagged strains were then transformed with pRS319 (RRID:Addgene_35459, ref. (79)) to introduce a *LEU3* marker for selection. In subsequent steps, diploid strains with HA-or Myc-tagged alleles were sporulated and haploid segregants were mated to yield the competition ChIP (CC) strains with different tags on each of the alleles for the GTF of interest. Proper integration and function of the targeted alleles were confirmed for all strains by PCR (Additional file 6: Table S5 for primers), Western blotting using anti-HA or anti-Myc antibodies, and targeted DNA sequencing of the modified loci.

### Western blotting

To measure the time course of synthesis of the *GAL1*-induced alleles, CC strains were grown in 175 ml YEP+2% raffinose. At OD600 of 0.6, a 20 ml aliquot of the culture was collected for the 0 min time point and 11 ml of 30% galactose was added to the remaining culture. 20 ml aliquots were removed at 10, 20, 25, 30, 40, 60, 90 and 120 minutes after galactose addition, and whole cell extracts were prepared from them as described previously (44). Whole cell extract protein was resolved on 10-12% SDS-Page gels (depending on the size of the tagged protein). The protein was transferred overnight to 0.22µ PVDF membranes and probed using either anti-HA (Abcam Cat# ab9110, RRID:AB_307019) or anti-Myc (Abcam Cat# ab32, RRID:AB_303599) antibodies followed by detection using either the HRP-conjugated goat anti-mouse secondary antibody, (for Myc; Thermo Fisher Scientific Cat# 31430, RRID:AB_228307) or goat anti-rabbit secondary antibody (for HA; Thermo Fisher Scientific Cat# 31460, RRID:AB_228341) and ECL substrate (Thermo Fisher Scientific Cat# 32106).

### CC time course experiments and ChIP-seq library preparation

Each CC strain was inoculated in 100 ml YEP+2% raffinose at 30° C and incubated overnight. These starter cultures were then used the next day to inoculate 2,250 ml cultures of YEP+2% raffinose at an initial OD600 of 0.05. When an OD of 0.6 was reached, for the 0 minute timepoint 250 ml of the culture was crosslinked by adding 6.75 ml formaldehyde (Thermo Fisher Scientific Cat# F79-500) to achieve a final concentration of 1% for 20 minutes. The reaction was then quenched by adding 15 ml of 2.5 M glycine for 5 minutes and the cells were collected by centrifugation. To the rest of the 2,000 ml culture, 142.8 ml of 30% galactose was added to yield a final concentration of 2%. At 10, 20, 25, 30, 40, 60, 90, and 120 minute time points, 250 ml of the culture was collected, crosslinked, and quenched the same way as the 0 min time point. Cell pellets were washed 3 times with TBS buffer (40 mM Tris-HCl, pH 7.5 plus 300 mM NaCl) and ChIP was performed as described (80). The HA and Myc antibodies used for ChIP were the same as those used for western blotting described above. Successful ChIP was confirmed by RT-PCR using primers to detect binding to the *URA3* promoter (5’-AAGATGCCCATCACCAAAA-3’ and 5’-AAGAATACCGGTTCCCGATG-3’). ChIP-seq libraries were prepared following the manufacturer’s instructions using the Illumina TruSeq ChIP library prep kit set A and B (Cat# IP-202-1012 and IP-202-1024). Successful amplification was confirmed by RT-PCR using the *URA3* promoter primers. Library quality was assessed using an Agilent Bioanalyzer 2100 and the Agilent-1000 DNA kit (Agilent Cat# 5067-1504), and libraries were quantified using the Qubit dsDNA Quantitation, High Sensitivity kit (Cat# Q32851). A 5nM pool of each library was sequenced on by the UVA Genome Analysis and Technology Core (RRID:SCR_018883) using Illumina NextSeq500 and NextSeq2000 instruments.

### Nascent RNA labelling

Nascent RNA labelling was performed as previously described (81). Briefly, W303 cells were grown as for competition ChIP and induced with 2% galactose for 20 or 60 minutes. An 800 ml culture in YEP +2% raffinose was grown at 30° C to an OD600 of 0.6, then 57ml of 30% galactose was added. Twenty minutes after galactose addition, 400 ml of the culture was divided into 200 ml aliquots and 500 μl of 2M 4-thiouracil (4-sU, Sigma-Aldrich Cat# 440736-1G) was added to one of the flasks with vigorous mixing and returned to the shaking incubator for 6 minutes. Cells with and without 4-sU were pelleted and washed with TBS. At the 60 minute time point the remaining 400 ml culture was split and treated as described for the 20 minute timepoint culture. Two biological replicates were obtained for each condition.

*S. pombe* strain SY78 cells were used as a spike-in normalization control. 100 ml of *S. pombe* cells were grown in YE media (0.5% yeast extract plus 3% glucose) to an OD600 of 0.6 and labelled by adding 125 μl of 2M 4-sU for 6 minutes and collected by centrifugation.

The *S. cerevisiae* W303 cells and *S. pombe* SY78 cells were mixed in an 8:1 ratio for each condition and RNA was isolated using the Ribopure Yeast Kit (Ambion Cat# AM1924). 40 μg of RNA was biotinylated with 4 μg of MTSEA Biotin XX (Biotium Cat# 90066). The biotinylated RNA was isolated by binding to 80 μl of a Dynabeads MyOne Streptavidin C1 bead suspension (Invitrogen Cat# 65001) by rotating the tube for 15 minutes, and the unbound supernatant was saved. The bound RNA was eluted in 50 μl of streptavidin elution buffer. The eluted RNA and the RNA in the flowthrough were purified and concentrated using RNeasy columns (Qiagen Cat# 74104).

### RNA-seq

Ribosomal RNA was depleted using the Ribo Minus Yeast module (Thermo Fisher Scientific Cat# 45-7013) and libraries were constructed using the Ultra Directional RNA Library Prep Kit (NEBNext Cat# E74205) and Multiplex Oligos (NEBNext Cat# E73355). Sequencing was performed by Novogene using the Illumina NovaSeq 6000 platform.

### Preprocessing of high throughput DNA sequencing data

Libraries prepared from each time point for a given GTF and for either HA-or Myc-tagged samples were sequenced in a single multiplexed run. Raw read quality was assessed using FASTQC (v0.11.5) (82). Fastq files from individual flow cells were merged and reads were mapped to the sacSer3 reference genome using Bowtie2 (v2.2.6) (83) with default settings. Overall read mapping was typically in the 90+% range, yielding ∼20-30M reads per time point on average. The resulting SAM files were converted to BAM format, unmapped reads were removed and the BAM files were sorted and indexed using SAMtools (v0.1.19-44428cd) (84). The landscape of read mapping was inspected using the Integrated Genomics Viewer (IGV) (85) and peaks of enrichment were identified using MACS2 (v2.1.0.20151222) (86) applied to each of several early time point Myc datasets with an input dataset as control and options --nomodel –extsize 147. Peaks from individual MACS2 runs were browsed in IGV, then concatenated and merged using the bedtools (v2.18.2) *merge* function (87). Count tables were then generated by associating reads with the peak intervals using bedtools *multicov*. Read counts were normalized in a three-step process. First, read counts in each peak and for each time point were normalized to the overall read depth. Next, read counts for the HA samples were normalized to the average relative levels of the factor of interest using the average values obtained from three independent western blots. Lastly, the normalized HA read count matrix was divided by the normalized Myc count matrix to yield the ratio count tables for mathematical modeling as described below. Importantly, this normalization approach was validated by comparison with earlier results: residence times derived from normalized TBP CC data were strikingly well correlated with TBP CC data obtained many years earlier and using arrays rather than sequencing (Additional file 1: Fig. S10c).

### Deriving residence times from competition ChIP-seq ratio data using a mass action kinetics turnover model

We adapted the approach of Zaidi et al. (36) originally developed for TBP competition ChIP-chip data, to fit a differential equation based turnover model at every GTF site using normalized competition ChIP-seq data from multiple GTFs. We used normalized count tables (see previous section of Methods) with HA/Myc ratios for every GTF site, *R*(*t*), for every timepoint, *t*. We ultimately estimate the ratio of fractional occupancies of HA-over Myc-tagged GTF, θ_*B*_(*t*)/θ_*A*_(*t*) with *B* and *A* representing HA- and Myc-tagged proteins, respectively, from *R*(*t*) at every timepoint. We then fit a mass action kinetic turnover model to the estimated ratio of fractional occupancies at every promoter site where a peak was identified. More specifically, we first fit the normalized ratio of HA-over Myc-tagged relative protein levels as estimated by Western blotting versus induction time, which we denote *c*_*B*_(*t*)/*c*_*A*_with *B* and *A* representing HA- and Myc-tagged protein, respectively, to a Hill model

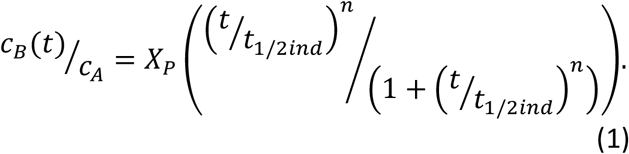

In Additional file 7: Table S6, we show the resulting fitted parameters (*X*_*P*_, *t*_1/2*ind*_) and statistics associated with the significance of each parameter’s contribution to the fit for every GTF. In this case, we fixed the Hill coefficient, *n*, to be an integer and selected the value that maximized the adjusted *R*^2^. In order to satisfy the *t* = 0 and *t* → ∞ boundary condition of the mass action kinetic turnover model shown below in Eqs. (2) and (3), which are θ_*B*_(0)/θ_*A*_(0) = 0 and 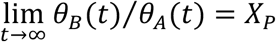 subtract the residual background and scale the normalized competition ChIP-seq ratio data at every site where peaks were called as follows. We fit the data to a Hill model with the form shown in Eq. (1) with the same *n* and an added background variable *B* at every site. This yields an amplitude, *X*_*CC*_, a half time rise, *t*_1/2*CC*_ and background *B* for every site. We estimate the ratio of HA-over Myc-tagged GTF occupancy, θ_*B*_(*t*)/θ_*A*_(*t*), at every site for every timepoint, *t*, by subtracting the residual background *B* from the normalized ChIP signal ratio data, *R*(*t*), and scaling the result: θ_*B*_(*t*)/θ_*A*_(*t*) = (*X*_*p*_/*X*_*CC*_) (*R*(*t*) −*B*). We then effectively solve the following coupled differential equations, which model each GTF’s turnover at every site which we assume follows mass action kinetics, where *k*_*a*_ and *k*_*d*_ are the molecular on- and off-rate respectively:

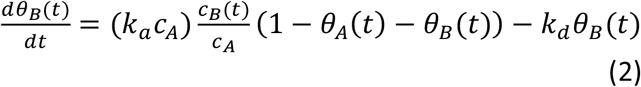

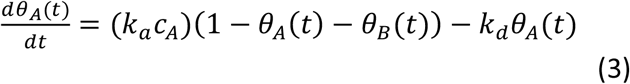

We assume that these rates are the same for both HA- and Myc-tagged GTFs. These coupled equations cannot be solved analytically. Thus, we effectively solve them and fit the resulting ratio of occupancies, θ_*B*_(*t*)/θ_*A*_(*t*), to the background subtracted, scaled competition ChIP-seq data using Mathematica. Briefly, we use the function *ParametricNDSolveValue* twice to return an effective, numerical solution of Eqs. (2) and (3) as a function of the parameters *k*_*a*_*c*_*A*_ and *k*_*d*_: θ_*B*_(*t*; *k*_*a*_*c*_*A*_, *k*_*d*_) and θ_*A*_(*t*; *k*_*a*_*c*_*A*_, *k*_*d*_). We then take the ratio of the outputs of ParametricNDSolveValue, θ_*B*_(*t*; *k*_*a*_*c*_*A*_, *k*_*d*_)/θ_*A*_(*t*; *k*_*a*_*c*_*A*_, *k*_*d*_), and input it into *NonlinearModelFit* which then fits this ratio to the background subtracted, scaled competition ChIP-seq data. We and others formally show the ratio of fractional occupancies is relatively insensitive to the on-rate, *k*_*a*_*c*_*A*_, while being highly sensitive to the offrate, *k*_*d*_. We derive the physical residence time for every GTF at every site using *t*_1/2_ = In 2 /*k*_d_. Finally, we make use of an observation made in (36) to make precise starting estimates of the residence time for non-linear model fitting using *NonlinearModelFit*. Specifically, the residence time is well approximated by a relatively simple linear or quadratic function of *t*_1/2*CC*_ − *t*_1/2*ind*_ derived by fitting a Hill model to the normalized competition ChIP-seq ratio data at every site and the ratio of GTF protein levels as a function of time. We start with an initial guess that works well for most GTFs: 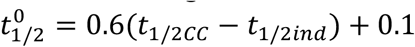 (Fig. 1d), perform the fit of the actual turnover model to the scaled, background subtracted competition ChIP-seq data, derive estimates of *t*_1/2_, fit *t*_1/2_ to linear or quadratic functions of 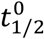 use this more precise relationship of initial estimate of residence time, *t*_1/2*CC*_ − *t*_1/2*ind*_, and refit the turnover model to the competition ChIP data. In Additional file 8: Table S7, we show the initialization formulas used for the final turnover model fit to the competition ChIP-seq data used to derive the final estimates of residence times for every GTF. Finally, *NonlinerModelFit* returns a number of statistics associated with the fit at every site. This includes an error estimate of the off-rate, ∆*k*_*d*_, and the adjusted *R*^2^. Sites that yielded a relative error ∆*k*_*d*_/*k*_*d*_ < 3 and adjusted *R*^2^ > 0.7 were used in downstream analysis involving residence time estimates.

### Fitting additional reliably fast sites

After initial fitting, additional reliably fast sites were added to the estimated residence times. These were identified through fitting Hill equation Eq. (1) with the R *nls* function to the normalized HA/Myc count ratios which were further normalized to range between zero and one. Hill coefficients were provided from protein induction curve fits (Fig. 2c, Additional file 1: Fig. S1e).

Initial estimates for fitting the Hill model using the *nls* function were set with parameter *start = list(t*_*1/2CC*_*=40, X*_*CC*_ *= 1)*, and parameter *control* was set to *nlc*. For each GTF, sites without estimated residence times from the turnover model whose Δ*t*_1/2_ = *t*_1/2*CC*_ − *t*_1/2*ind*_ (Fig. 1d) were less than 2 min were classified as reliably fast (< 1 min). All residence time estimates are available in Additional file 3: Table S2.

For plotting purposes, the residence times for the reliably fast sites were generated with the R *runif* function with *min=0, max=1*. At the beginning of each script, the function *set*.*seed* was used with parameter *42* for reproducibility. In each plot, the randomly generated values are highlighted either by their separation by dashed line or shaded area.

### Gene assignment and filtering

Individual regions were assigned to the nearest genes with *calcFeatureDist_aY* function (available from https://github.com/AubleLab/annotateYeast) with default parameters. Only regions within -250 to 100 bp from transcription start sites (TSSs) were kept. If multiple regions were assigned to one gene, only the closest one was kept. Regions assigned to tRNAs were removed from the analysis.

### Nascent RNA-seq analysis

Raw paired-end FASTQ files were mapped to the *S. cerevisiae* genome (http://daehwankimlab.github.io/hisat2/download/#s-cerevisiae, R64-1-1) with HISAT2 (2.0.4) (88) with parameter *--rna-strandness RF* and converted to BAM files using SAMtools (0.1.19-44428cd) (84) *view* function with parameters *-S -b*. SAMtools *sort* and *index* functions with default parameters were used to sort and index the BAM alignment files.

To create alignment indexes for *S. pombe* (used for normalization), the *S. pombe* FASTA file (ASM294v2) was obtained from Ensembl (89) and converted to an index files with the *hisat2-build* function with default parameters. The paired-end FASTQ files were then mapped against the created index files and further processed analogously to *S. cerevisiae*.

The quality of both FASTQ and BAM files was assessed with FastQC (0.11.5) (82) in combination with multiQC (v1.11) (90) and BAM files were further visually inspected with IGV (2.7.2) (85).

The aligned reads were quantified over *S. cerevisiae* genes using Rsubread (2.4.3) (91) *featureCounts* function with parameters *GTF*.*featureType=“gene”, GTF*.*attrType=“gene_id”, countMultiMappingReads=TRUE, strandSpecific=2, isPairedEnd=TRUE*. The GTF and FASTA files provided to the function were obtained from Ensembl (89), genome assembly R64-1-1. To normalize the data, normalization factors for each sample were calculated as the total number of reads mapped to *S. pombe* divided by 2,000,000. The normalized counts were obtained by dividing the raw counts by each sample’s corresponding normalization factor. Genes with 0 counts in more than half of the samples were filtered out.

Principal component analysis (PCA) was performed by first creating a *DESeq* object from the raw count table (with low count genes filtered out) with the DESeq2 (1.30.1) (92) *DESeqDataSetFromMatrix* function followed by *S. pombe* normalization with DESeq2 *normalizationFactors* and regularized log transformation with DESeq2 *rlog* function with parameter *blind=TRUE*. The resulting object was passed to R *prcomp* function.

DESeq2 was used to identify any differences in gene expression between samples grown for 20 or 60 minutes in galactose. Raw counts from samples with thiouracil addition were passed to *DESeqDataSetFromMatrix* function with *design* parameter set to *time in galactose*. S. pombe normalization factors were set with *normalizationFactors*. Genes with adjusted p-value (padj) < 0.05 were considered differentially expressed between the two conditions.

Synthesis rates were estimated with DTA (2.36.0) (93) *DTA*.*estimate* function. *S. pombe*-normalized counts from samples with thiouracil addition were used for the analysis. All genes with 0 count in any of the samples were filtered out and the final matrix passed to the function. All genes from the final filtered matrix were passed to the parameter *reliable*. Further parameters were set to: *tnumber=Sc*.*tnumber, check=TRUE, ccl = 150, mRNAs=60000, condition=“real_data”, ratiomethod=“bias”*, and time in the *phenomat* object was set to *6*. Final synthesis rates in mRNA per cell per minute were obtained by dividing the synthesis rates output from the *DTA*.*estimate* function by 150 (length of the cell cycle in minutes). The final synthesis rates are available in Additional file 4: Table S3. Comparison of synthesis rates between samples grown for 20 vs. 60 minutes in galactose was performed using *DTA*.*dynamic*.*estimate* function similarly as described above with additional columns *timeframe* and *timecourse* in the *phenomat* object specifying 20 vs. 60 minute conditions. The correlation between the synthesis rates of the two time courses was calculated using the R *cor* function with *method=“pearson”*.

### Comparison with other data

TBP residence time estimates were obtained from Zaidi et al, 2017 (36), TBP and TFIIE residence time estimates from Zaidi, Hoffman et al, 2017 (44), transcription rates from Garciá-Martinez et al, 2004 (38), and Rap1 residence times from Lickwar et al, 2012 (40). Correlations were calculated with R *cor* function. For residence time correlations, where we do not have exact time estimates for fast sites, Pearson’s correlation was used, while for synthesis rates, Spearman’s rank correlation was used.

### Model plotting

Examples of model fits were obtained by extracting Hill equation coefficients, as described in “Fitting additional reliably fast sites” section of Methods. Output model values and the measured competition ChIP (CC) values were both scaled to range between zero and one to create comparable plots by dividing the values by the estimated *X*_*cc*_ parameter.

### Visual inspection with genome browser

To view the normalized HA/Myc ratios in the genome browser, BAM alignment files were first converted to bigWig files using the deepTools (3.3.1) (94) *bamCoverage* function. The parameter *scaleFactor* was set to per million mapped reads scaling factor for the Myc samples and to per million mapped reads multiplied by HA/Myc protein induction ratio for the HA samples. The final log_2_ transformed ratios of HA/Myc were obtained by passing the generated bigWig files to the deepTools *bigwigCompare* function with parameter *operation* set to *log2*.

### Residence time vs. synthesis rate

To explore the residence times of each analyzed GTF in relationship to synthesis rates, synthesis rates were first divided into quartiles using the R *ntile* function with the parameter *ngroups* set to 4. Residence times within each synthesis quartile were plotted as boxplots with ggplot2 (3.3.6) (95) *geom_boxplot* function, where the middle line represents the median, the lower and upper hinge represent the first and third quartiles, and the whiskers represent 1.5 * interquartile range of the values.

The correlations between synthesis rates and residence times were calculated with R *cor* function with *method* set to *“pearson”*.

Linear models between synthesis rates were built with R *lm* function either as linear models between synthesis rate and residence times of individual GTFs or as a linear model between synthesis rates and a linear combination of residence times of all factors in one model.

### Transcription efficiency

Transcription efficiency (TE) was obtained by multiplying the synthesis rate by residence time of a given TF. The log_2_ transformed values were plotted with ggplot2 *geom_violin* function to better represent the efficiency of a binding event to produce an RNA molecule (values below zero represent multiple binding events for RNA molecule synthesis). Medians of the log2 transformed TE values for each TF were added to the violin plots with the tidyverse (1.3.1) (96) *stat_summary* function with parameter *fun=median*.

### PCA

To represent genes or GTFs using their corresponding high dimensional data in low dimensional space, we performed PCA on the residence times with or without exclusion of the reliably fast sites. Since the residence time estimates for all TFs were not available for all genes the missing values were imputed with the missMDA (1.18) package (97). The table containing the reliable residence times was first passed to the *estim_ncpPCA* function with parameter *method*.*cv* set to *“Kfold”*. The residence time table was then passed to the *imputePCA* function along with the *ncp* object outputted from the *estim_ncpPCA* function. The c*ompleteObs* object from the outputted list was then passed to the *prcomp* function with parameter *scale*.*=TRUE* to obtain the principal components. Depending on the orientation of the input matrix passed to the *prcomp* function, principal components representing genes or GTFs were obtained. To color-code the PCA plot with mean synthesis rates, the tidyverse (1.3.1) (96) function *stat_summary_2d* was used with parameter *z* set to the synthesis rates and parameter color set to *“transparent”*. Viridis (0.6.2) (98) color scale *“B”* was used for coloring. The first two principal components from the “gene-oriented” PCA matrices were then correlated with the residence times of each TF and with the synthesis rates using the R function *cor* with *method=“pearson”*.

### Residence time and synthesis rate comparison between gene classes

The list of genes with TATA-containing promoters was obtained From Rhee and Pugh, 2012 (99). Genes were classified as ribosomal subunit if their systematic name started with “RPL”. To compare the residence times and synthesis rates between classes, a two-tailed t-test was carried out with results plotted using the ggpubr (0.4.0) (100) *stat_compare_means* function with parameters set to *method = “t*.*test”, label = “p*.*signif”*. The symbols indicate the following: n.s. p > 0.05, * p <= 0.05, ** p <= 0.01, *** p <= 0.001, and **** p <= 0.0001. To compare residence times across synthesis quartiles, synthesis rates were separated into the four quartiles based on synthesis rates within each group (e.g. TATA-containing and TATA-less). Box plots were created using ggplot2 (3.3.6) (95) *geom_boxplot* function, where the middle line represents the median, the lower and upper hinge represent the first and third quartiles, and the whiskers represent 1.5 * interquartile range of the values.

### Heatmap

Only genes for which residence times were available across all GTFs (except for excluded TBP, whose residence times are mostly <1 minute and would therefore present mostly randomly generated values) were included in the heatmap (n = 1417). Reliable fast residence times were replaced by randomly generated values between zero and one (function *runif: min=0, max=1*; *set*.*seed(42)*). Prior to plotting, residence times for each factor were z-score normalized using the R function *scale* with default settings. A final heatmap was created with the ComplexHeatmap (2.6.2) (101) function *Heatmap* with parameters set to *clustering_method_rows=“ward*.*D”, row_split=10*. Genes belonging to each of the 10 clusters (Additional file 5: Table S4) were extracted from the heatmap object and mean synthesis rates for each cluster were calculated.

### Functional annotation

Genes belonging to each heatmap cluster were passed to g:Profiler (102) for pathway enrichment. In g:Profiler, *S. cerevisiae* S88C was selected as organism and data sources were set to GO molecular function (GO:MF), GO biological process (GO:BP), KEGG, WikiPathways (WP), and TRANSFAC. Additionally, genes from the clusters were tested for enrichment within genes associated with DNA-binding factors (DBFs) from Rossi et al, 2021 (13), here referred to as Yeast Epigenome database (see section “Yeast DBF database (Yeast Epigenime)” of the Methods for information about data accessions and curation). Enrichment was established by performing Fisher’s exact test (R function *fisher*.*test*, parameter *alternative=“greater”*), where the universe was set to the union of all genes involved in the heatmap and all genes associated with a given factor. Final p-values were corrected for multiple testing with false discovery rate (FDR, R function *p*.*adjust*: *method=“fdr”*). Results with FDR padj < 0.05 or p < 0.05 were considered significant.

### Yeast DBF database (Yeast Epigenome)

BED files from Rossi et al, 2021 (13) were obtained from Gene Expression Omnibus under accession number GSE147927. Replicates were merged with the bedtools (v2.29.2) (87) *merge* function after they were sorted with the base Linux *sort* function with parameters *-k1,1 -k2,2n*. Regions were then assigned to genes analogously to assignment of the CC regions (see “Gene assignment and filtering” section of Methods). The output consists of gene lists for individual DBFs within promoter regions.

### Additional tools used

Tidyverse (1.3.1) package (96) was used for data processing in R, ggplot2 (3.3.6) (95) was used for plotting. Illustrations were made with Biorender (https://biorender.com/). Figures were assembled with Inkscape (1.0.2, https://inkscape.org/).

## Abbreviations

CC: Competition ChIP
DBF: DNA binding factor
DTA: dynamic transcriptome analysis
FDR: false discovery rate
GTF: general transcription factor
GO:BP: gene ontology biological process
GO:MF: gene ontology molecular function
GTF: general transcription factor
padj: adjusted p-value
PC1: first principal component
PC2: second principal component
PCA: principal component analysis
PIC: preinitiation complex
Pol II: RNA polymerase II
TBP: TATA-binding protein
TE: transcription efficiency
TF: transcription factor
TSS: transcription start site
WP: WikiPathways

## Declarations

### Ethics approval and consent to participate

Not applicable.

### Consent for publication

Not applicable.

### Availability of data and materials

The datasets supporting the conclusions of this article are available in the GEO repository, GSE235002. The CC samples are available from GSE235000, and nascent RNA-seq samples from GSE235001. The scripts are available from https://github.com/AubleLab/PIC_competition_ChIP_scripts (103).

## Competing interests

The authors declare that they have no competing interests.

## Funding

Funding for this work was provided by National Institutes of Health (Grant R01 GM55763 to DTA) and the Biomedical Sciences Graduate Program, University of Virginia (Wagner fellowship to KK). The funders did not have any role in the design of the study and collection, analysis, and interpretation of the data nor in the writing of the manuscript.

## Authors’ contributions

EAH and SJS performed the yeast molecular biological and molecular genetic manipulations. Samples and libraries for CC were obtained by SJS. Preprocessing of the data and initial analyses were performed by DTA. SB developed the kinetic model and performed kinetic analysis of the CC data using the mass-action model. KK analyzed the residence time results, analyzed the nascent RNA-seq data, and generated all of the figures. All authors contributed to writing the manuscript.

## Acknowledgements

We are grateful to Patrick Grant for discussions and critical reading of the manuscript.

## List of additional files

**Additional file 1**.

Supplementary figures (S1-S9), Supplementary references.

**Additional file 2**.

Table S1. Normalized quantified Western blots.

**Additional file 3**.

Table S2. Residence times.

**Additional file 4**.

Table S3. Synthesis rates.

**Additional file 5**.

Table S4. Heatmap clusters.

**Additional file 6**.

Table S5. List of primers.

**Additional file 7**.

Table S6. Hill model fits to HA/Myc Western blot data for turnover model.

**Additional file 8**.

Table S7. Accurate initial estimates of residence times for turnover model fits.

## Notes

https://www.ncbi.nlm.nih.gov/geo/query/acc.cgi?acc=GSE235002

https://www.ncbi.nlm.nih.gov/geo/query/acc.cgi?acc=GSE235000

https://www.ncbi.nlm.nih.gov/geo/query/acc.cgi?acc=GSE235001

### Competing Interest Statement

The authors have declared no competing interest.

## References

1. He Y, Yan C, Fang J, Inouye C, Tjian R, Ivanov I, et al. Near-atomic resolution visualization of human transcription promoter opening. Nature. 2016;533(7603):359–65.

2. Plaschka C, Hantsche M, Dienemann C, Burzinski C, Plitzko J, Cramer P. Transcription initiation complex structures elucidate DNA opening. Nature. 2016;533(7603):353–8.

3. Sainsbury S, Bernecky C, Cramer P. Structural basis of transcription initiation by RNA polymerase II. Nat Rev Mol Cell Biol. 2015;16(3):129–43.

4. Hahn S, Young ET. Transcriptional Regulation in Saccharomyces cerevisiae: Transcription Factor Regulation and Function, Mechanisms of Initiation, and Roles of Activators and Coactivators. Genetics. 2011 Nov 1;189(3):705–36.

5. Hahn S. Structure and mechanism of the RNA polymerase II transcription machinery. Nat Struct Mol Biol. 2004;11(5):394–403.

6. Nogales E, Louder RK, He Y. Structural Insights into the Eukaryotic Transcription Initiation Machinery. Annu Rev Biophys. 2017 May 22;46(1):59–83.

7. Patel AB, Louder RK, Greber BJ, Grünberg S, Luo J, Fang J, et al. Structure of human TFIID and mechanism of TBP loading onto promoter DNA. Science (80-). 2018 Dec 21;362(6421).

8. Ehara H, Yokoyama T, Shigematsu H, Yokoyama S, Shirouzu M, Sekine SI. Structure of the complete elongation complex of RNA polymerase II with basal factors. Science (80-). 2017 Sep 1;357(6354):921–4.

9. Hahn S, Buratowski S. Snapshots of transcription initiation. Nature. 2016;533(7603):331–2.

10. Venters BJ, Wachi S, Mavrich TN, Andersen BE, Jena P, Sinnamon AJ, et al. A Comprehensive Genomic Binding Map of Gene and Chromatin Regulatory Proteins in Saccharomyces. Mol Cell. 2011 Feb 18;41(4):480–92.

11. Harbison CT, Gordon DB, Lee TI, Rinaldi NJ, Macisaac KD, Danford TW, et al. Transcriptional regulatory code of a eukaryotic genome. Nature. 2004;431(7004):99–104.

12. Yen K, Vinayachandran V, Batta K, Koerber RT, Pugh BF. Genome-wide Nucleosome Specificity and Directionality of Chromatin Remodelers. Cell. 2012 Jun 22;149(7):1461–73.

13. Rossi MJ, Kuntala PK, Lai WKM, Yamada N, Badjatia N, Mittal C, et al. A high-resolution protein architecture of the budding yeast genome. Nat 2021 5927853. 2021 Mar 10;592(7853):309–14.

14. Patel AB, Greber BJ, Nogales E. Recent insights into the structure of TFIID, its assembly, and its binding to core promoter. Curr Opin Struct Biol. 2020 Apr 1;61:17–24.

15. Louder RK, He Y, López-Blanco JR, Fang J, Chacón P, Nogales E. Structure of promoter-bound TFIID and model of human pre-initiation complex assembly. Nature. 2016;531(7596):604–9.

16. Chen H, Pugh BF. What do Transcription Factors Interact With? J Mol Biol. 2021;433(14):166883.

17. Farnung L, Vos SM. Assembly of RNA polymerase II transcription initiation complexes. Curr Opin Struct Biol. 2022;73:102335.

18. Tsutakawa SE, Tsai C-L, Yan C, Bralić A, Chazin WJ, Hamdan SM, et al. Envisioning how the prototypic molecular machine TFIIH functions in transcription initiation and DNA repair. DNA Repair (Amst). 2020;96:102972.

19. Rimel JK, Taatjes DJ. The essential and multifunctional TFIIH complex. Protein Sci. 2018 Jun 1;27(6):1018–37.

20. Schilbach S, Hantsche M, Tegunov D, Dienemann C, Wigge C, Urlaub H, et al. Structures of transcription pre-initiation complex with TFIIH and Mediator. Nature. 2017;551(7679):204–9.

21. Malik S, Molina H, Xue Z. PIC Activation through Functional Interplay between Mediator and TFIIH. J Mol Biol. 2017;429(1):48–63.

22. Nguyen VQ, Ranjan A, Liu S, Tang X, Ling YH, Wisniewski J, et al. Spatiotemporal coordination of transcription preinitiation complex assembly in live cells. Mol Cell. 2021;81(17):3560-3575.e6.

23. Guglielmi B, LaRochelle N, Tjian R. Gene-Specific Transcriptional Mechanisms at the Histone Gene Cluster Revealed by Single-Cell Imaging. Mol Cell. 2013 Aug 22;51(4):480–92.

24. Luse DS. The RNA polymerase II preinitiation complex. Transcription. 2014;5(1):e27050.

25. Sikorski TW, Buratowski S. The basal initiation machinery: beyond the general transcription factors. Curr Opin Cell Biol. 2009 Jun 1;21(3):344–51.

26. Sanchez A, Golding I. Genetic determinants and cellular constraints in noisy gene expression. Science (80-). 2013 Dec 6;342(6163):1188–93.

27. Raser JM, O’Shea EK. Control of stochasticity in eukaryotic gene expression. Science (80-). 2004 Jun 18;304(5678):1811–4.

28. 28. Boeger H, Griesenbeck J, Kornberg RD. Nucleosome Retention and the Stochastic Nature of Promoter Chromatin Remodeling for Transcription. Cell. 2008 May 16;133(4):716–26.

29. Brown CR, Boeger H. Nucleosomal promoter variation generates Gene expression noise. Proc Natl Acad Sci U S A. 2014 Dec 16;111(50):17893–8.

30. Lenstra TL, Rodriguez J, Chen H, Larson DR. Transcription Dynamics in Living Cells. Annu Rev Biophys. 2016 Jul 5;45(1):25–47.

31. Boettiger AN, Ralph PL, Evans SN. Transcriptional Regulation: Effects of Promoter Proximal Pausing on Speed, Synchrony and Reliability. PLOS Comput Biol. 2011;7(5):e1001136.

32. Ravarani CNJ, Chalancon G, Breker M, de Groot NS, Babu MM. Affinity and competition for TBP are molecular determinants of gene expression noise. Nat Commun. 2016;7(1):10417.

33. Hager GL, McNally JG, Misteli T. Transcription Dynamics. Mol Cell. 2009 Sep 24;35(6):741–53.

34. Voss TC, Hager GL. Dynamic regulation of transcriptional states by chromatin and transcription factors. Nat Rev Genet. 2014;15(2):69–81.

35. Lickwar CR, Mueller F, Lieb JD. Genome-wide measurement of protein-DNA binding dynamics using competition ChIP. Nat Protoc. 2013;8(7):1337–53.

36. Zaidi HA, Auble DT, Bekiranov S. RNA synthesis is associated with multiple TBP-chromatin binding events. Sci Reports 2017 71. 2017 Jan 4;7(1):1–12.

37. Miller C, Schwalb B, Maier K, Schulz D, Dümcke S, Zacher B, et al. Dynamic transcriptome analysis measures rates of mRNA synthesis and decay in yeast. Mol Syst Biol. 2011;

38. García-Martínez J, Aranda A, Pérez-Ortín JE. Genomic Run-On Evaluates Transcription Rates for All Yeast Genes and Identifies Gene Regulatory Mechanisms. Mol Cell. 2004 Jul 23;15(2):303–13.

39. Basehoar AD, Zanton SJ, Pugh BF. Identification and Distinct Regulation of Yeast TATA Box-Containing Genes. Cell. 2004 Mar 5;116(5):699–709.

40. Lickwar CR, Mueller F, Hanlon SE, McNally JG, Lieb JD. Genome-wide protein–DNA binding dynamics suggest a molecular clutch for transcription factor function. Nat 2012 4847393. 2012 Apr 11;484(7393):251–5.

41. van Werven FJ, van Teeffelen HAAM, Holstege FCP, Timmers HTM. Distinct promoter dynamics of the basal transcription factor TBP across the yeast genome. Nat Struct Mol Biol. 2009;16(10):1043–8.

42. Hasegawa Y, Struhl K. Promoter-specific dynamics of TATA-binding protein association with the human genome. Genome Res. 2019 Dec 1;29(12):1939–50.

43. Poorey K, Viswanathan R, Carver MN, Karpova TS, Cirimotich SM, McNally JG, et al. Measuring chromatin interaction dynamics on the second time scale at single-copy genes. Science (80-). 2013 Oct 18;342(6156):369–72.

44. Zaidi HA, Hoffman EA, Shetty SJ, Bekiranov S, Auble DT. Second-generation method for analysis of chromatin binding with formaldehyde–cross-linking kinetics. J Biol Chem. 2017 Nov 24;292(47):19338–55.

45. van Royen ME, Zotter A, Ibrahim SM, Geverts B, Houtsmuller AB. Nuclear proteins: finding and binding target sites in chromatin. Chromosom Res. 2011;19(1):83–98.

46. Paakinaho V, Presman DM, Ball DA, Johnson TA, Schiltz RL, Levitt P, et al. Single-molecule analysis of steroid receptor and cofactor action in living cells. Nat Commun. 2017;8(1):15896.

47. Normanno D, Dahan M, Darzacq X. Intranuclear mobility and target search mechanisms of transcription factors: A single-molecule perspective on gene expression. Biochim Biophys Acta - Gene Regul Mech. 2012;1819(6):482–93.

48. Swinstead EE, Miranda TB, Paakinaho V, Baek S, Goldstein I, Hawkins M, et al. Steroid Receptors Reprogram FoxA1 Occupancy through Dynamic Chromatin Transitions. Cell. 2016;165(3):593–605.

49. Liu Z, Tjian R. Visualizing transcription factor dynamics in living cells. J Cell Biol. 2018;217(4):1181–91.

50. Lionnet T, Wu C. Single-molecule tracking of transcription protein dynamics in living cells: seeing is believing, but what are we seeing? Curr Opin Genet Dev. 2021;67:94–102.

51. Brouwer I, Lenstra TL. Visualizing transcription: key to understanding gene expression dynamics. Curr Opin Chem Biol. 2019;51:122–9.

52. Zhang Z, English BP, Grimm JB, Kazane SA, Hu W, Tsai A, et al. Rapid dynamics of general transcription factor TFIIB binding during preinitiation complex assembly revealed by single-molecule analysis. Genes Dev. 2016 Sep 15;30(18):2106–18.

53. Sprouse RO, Karpova TS, Mueller F, Dasgupta A, McNally JG, Auble DT. Regulation of TATA-binding protein dynamics in living yeast cells. Proc Natl Acad Sci. 2008 Sep;105(36):13304–8.

54. Donovan BT, Huynh A, Ball DA, Patel HP, Poirier MG, Larson DR, et al. Live-cell imaging reveals the interplay between transcription factors, nucleosomes, and bursting. EMBO J. 2019 Jun 17;38(12):e100809.

55. Coulon A, Chow CC, Singer RH, Larson DR. Eukaryotic transcriptional dynamics: from single molecules to cell populations. Nat Rev Genet [Internet]. 2013;14(8):572–84. Available from: https://doi.org/10.1038/nrg3484

56. Nicolas D, Phillips NE, Naef F. What shapes eukaryotic transcriptional bursting? Mol BioSyst. 2017;13(7):1280–90.

57. Hipp L, Beer J, Kuchler O, Reisser M, Sinske D, Michaelis J, et al. Single-molecule imaging of the transcription factor SRF reveals prolonged chromatin-binding kinetics upon cell stimulation. Proc Natl Acad Sci. 2019 Jan 15;116(3):880–9.

58. Stone NR, Gifford CA, Thomas R, Pratt KJB, Samse-Knapp K, Mohamed TMA, et al. Context-Specific Transcription Factor Functions Regulate Epigenomic and Transcriptional Dynamics during Cardiac Reprogramming. Cell Stem Cell. 2019;25(1):87-102.e9.

59. Mony VK, Drangowska-Way A, Albert R, Harrison E, Ghaddar A, Horak MK, et al. Context-specific regulation of lysosomal lipolysis through network-level diverting of transcription factor interactions. Proc Natl Acad Sci. 2021 Oct 12;118(41):e2104832118.

60. Fertig EJ, Favorov A V, Ochs* MF. Identifying Context-Specific Transcription Factor Targets From Prior Knowledge and Gene Expression Data. IEEE Trans Nanobioscience. 2013;12(3):142–9.

61. Osman S, Cramer P. Structural Biology of RNA Polymerase II Transcription: 20 Years On. Annu Rev Cell Dev Biol [Internet]. 2020 Oct 6;36(1):1–34. Available from: https://doi.org/10.1146/annurev-cellbio-042020-021954

62. Bushnell DA, Bamdad C, Kornberg RD. A Minimal Set of RNA Polymerase II Transcription Protein Interactions. J Biol Chem. 1996 Aug 16;271(33):20170–4.

63. Ranish JA, Hahn S. Transcription: basal factors and activation. Curr Opin Genet Dev. 1996 Apr 1;6(2):151–8.

64. Orphanides G, Lagrange T, Reinberg D. The general transcription factors of RNA polymerase II. Genes Dev. 1996;10(21):2657–83.

65. Stasevich TJ, Hayashi-Takanaka Y, Sato Y, Maehara K, Ohkawa Y, Sakata-Sogawa K, et al. Regulation of RNA polymerase II activation by histone acetylation in single living cells. Nature [Internet]. 2014;516(7530):272–5. Available from: https://doi.org/10.1038/nature13714

66. Petrenko N, Jin Y, Dong L, Wong KH, Struhl K. Requirements for RNA polymerase II preinitiation complex formation in vivo. Green MR, Manley JL, editors. Elife. 2019;8:e43654.

67. Yudkovsky N, Ranish JA, Hahn S. A transcription reinitiation intermediate that is stabilized by activator. Nature. 2000;408(6809):225–9.

68. Tyree CM, George CP, Lira-De Vito LM, Wampler SL, Dahmus ME, Zawel L, et al. Identification of a minimal set of proteins that is sufficient for accurate initiation of transcription by RNA polymerase II. Genes Dev. 1993 Jul 1;7(7a):1254–65.

69. Fujiwara R, Murakami K. In vitro reconstitution of yeast RNA polymerase II transcription initiation with high efficiency. Methods. 2019;159–160:82–9.

70. Luse DS. Insight into promoter clearance by RNA polymerase II. Proc Natl Acad Sci. 2019 Nov 5;116(45):22426–8.

71. Greber BJ, Toso DB, Fang J, Nogales E. The complete structure of the human TFIIH core complex. Grigorieff N, Wolberger C, Grigorieff N, Darst SA, Berger JM, editors. Elife. 2019;8:e44771.

72. Nogales E, Greber BJ. High-resolution cryo-EM structures of TFIIH and their functional implications. Curr Opin Struct Biol. 2019;59:188–94.

73. Nozawa K, Schneider TR, Cramer P. Core Mediator structure at 3.4 Å extends model of transcription initiation complex. Nature. 2017;545(7653):248–51.

74. Plaschka C, Larivière L, Wenzeck L, Seizl M, Hemann M, Tegunov D, et al. Architecture of the RNA polymerase II–Mediator core initiation complex. Nature. 2015;518(7539):376–80.

75. Ralser M, Kuhl H, Ralser M, Werber M, Lehrach H, Breitenbach M, et al. The Saccharomyces cerevisiae W303-K6001 cross-platform genome sequence: insights into ancestry and physiology of a laboratory mutt. Open Biol. 2012 Aug 1;2(8):120093.

76. Longtine MS, Mckenzie III A, Demarini DJ, Shah NG, Wach A, Brachat A, et al. Additional modules for versatile and economical PCR-based gene deletion and modification in Saccharomyces cerevisiae. Yeast. 1998 Jul 1;14(10):953–61.

77. Gauss R, Trautwein M, Sommer T, Spang A. New modules for the repeated internal and N-terminal epitope tagging of genes in Saccharomyces cerevisiae. Yeast. 2005 Jan 15;22(1):1–12.

78. Güldener U, Heck S, Fiedler T, Beinhauer J, Hegemann JH. A New Efficient Gene Disruption Cassette for Repeated Use in Budding Yeast. Nucleic Acids Res. 1996 Jul 1;24(13):2519–24.

79. Sikorski RS, Hieter P. A system of shuttle vectors and yeast host strains designed for efficient manipulation of DNA in Saccharomyces cerevisiae. Genetics. 1989 May 1;122(1):19–27.

80. Viswanathan R, Hoffman EA, Shetty SJ, Bekiranov S, Auble DT. Analysis of chromatin binding dynamics using the crosslinking kinetics (CLK) method. Methods. 2014 Dec 1;70(2–3):97–107.

81. Warfield L, Ramachandran S, Baptista T, Devys D, Tora L, Hahn S. Transcription of Nearly All Yeast RNA Polymerase II-Transcribed Genes Is Dependent on Transcription Factor TFIID. Mol Cell. 2017;68(1):118-129.e5.

82. Andrews S. FastQC: a quality control tool for high throughput sequence data. [Internet]. 2010. Available from: https://www.bioinformatics.babraham.ac.uk/projects/fastqc/

83. Langmead B, Salzberg SL. Fast gapped-read alignment with Bowtie 2. Nat Methods. 2012;9(4):357–9.

84. Li H, Handsaker B, Wysoker A, Fennell T, Ruan J, Homer N, et al. The Sequence Alignment/Map format and SAMtools. Bioinformatics. 2009 Aug;25(16):2078–9.

85. Thorvaldsdottir H, Robinson JT, Mesirov JP. Integrative Genomics Viewer (IGV): high-performance genomics data visualization and exploration. Brief Bioinform. 2013 Mar 1;14(2):178–92.

86. Zhang Y, Liu T, Meyer CA, Eeckhoute J, Johnson DS, Bernstein BE, et al. Model-based analysis of ChIP-Seq (MACS). Genome Biol. 2008;9(9):R137.

87. Quinlan AR, Hall IM. BEDTools: a flexible suite of utilities for comparing genomic features. Bioinformatics. 2010 Mar 15;26(6):841–2.

88. Kim D, Paggi JM, Park C, Bennett C, Salzberg SL. Graph-based genome alignment and genotyping with HISAT2 and HISAT-genotype. Nat Biotechnol 2019 378. 2019 Aug 2;37(8):907–15.

89. Cunningham F, Allen JE, Allen J, Alvarez-Jarreta J, Amode MR, Armean IM, et al. Ensembl 2022. Nucleic Acids Res. 2022 Jan 7;50(D1):D988–95.

90. Ewels P, Magnusson M, Lundin S, Käller M. MultiQC: summarize analysis results for multiple tools and samples in a single report. Bioinformatics. 2016 Oct 1;32(19):3047–8.

91. Liao Y, Smyth GK, Shi W. featureCounts: an efficient general purpose program for assigning sequence reads to genomic features. Bioinformatics. 2014 Apr 1;30(7):923–30.

92. Love MI, Huber W, Anders S. Moderated estimation of fold change and dispersion for RNA-seq data with DESeq2. Genome Biol. 2014 Dec 5;15(12):1–21.

93. Schwalb B, Schulz D, Sun M, Zacher B, Dümcke S, Martin DE, et al. Measurement of genome-wide RNA synthesis and decay rates with Dynamic Transcriptome Analysis (DTA). Bioinformatics. 2012 Mar 15;28(6):884–5.

94. Ramírez F, Dündar F, Diehl S, Grüning BA, Manke T. DeepTools: A flexible platform for exploring deep-sequencing data. Nucleic Acids Res. 2014 Jul 1;42(W1):W187.

95. Wickham H. Ggplot2: Elegant graphics for data analysis. 2nd ed. Cham, Switzerland: Springer International Publishing; 2016.

96. Wickham H, Averick M, Bryan J, Chang W, McGowan L, François R, et al. Welcome to the Tidyverse. J Open Source Softw. 2019 Nov 21;4(43):1686.

97. Josse J, Husson F. missMDA: A Package for Handling Missing Values in Multivariate Data Analysis. J Stat Softw. 2016 Apr 4;70:1–31.

98. Garnier, Simon, Ross, Noam, Rudis, Robert, et al. viridis - Colorblind-Friendly Color Maps for R [Internet]. Available from: https://sjmgarnier.github.io/viridis/

99. Rhee HS, Pugh BF. Genome-wide structure and organization of eukaryotic pre-initiation complexes. Nat 2012 4837389. 2012 Jan 18;483(7389):295–301.

100. Kassambara A. ggpubr: “ggplot2” Based Publication Ready Plots. 2020.

101. Gu Z, Eils R, Schlesner M. Complex heatmaps reveal patterns and correlations in multidimensional genomic data. Bioinformatics. 2016 Sep 15;32(18):2847–9.

102. Raudvere U, Kolberg L, Kuzmin I, Arak T, Adler P, Peterson H, et al. G:Profiler: A web server for functional enrichment analysis and conversions of gene lists (2019 update). Nucleic Acids Res. 2019 Jul 1;47(W1):W191–8.

103. Kupkova K, Bekiranov S, Auble DT. PIC_competition_ChIP_scripts [Internet]. 2023. Available from: https://doi.org/10.5281/zenodo.8161712

