## Additional file 1 for "Genome-scale chromatin interaction dynamic measurements for key components of the RNA Pol II general transcription machinery"

### Supplementary figures

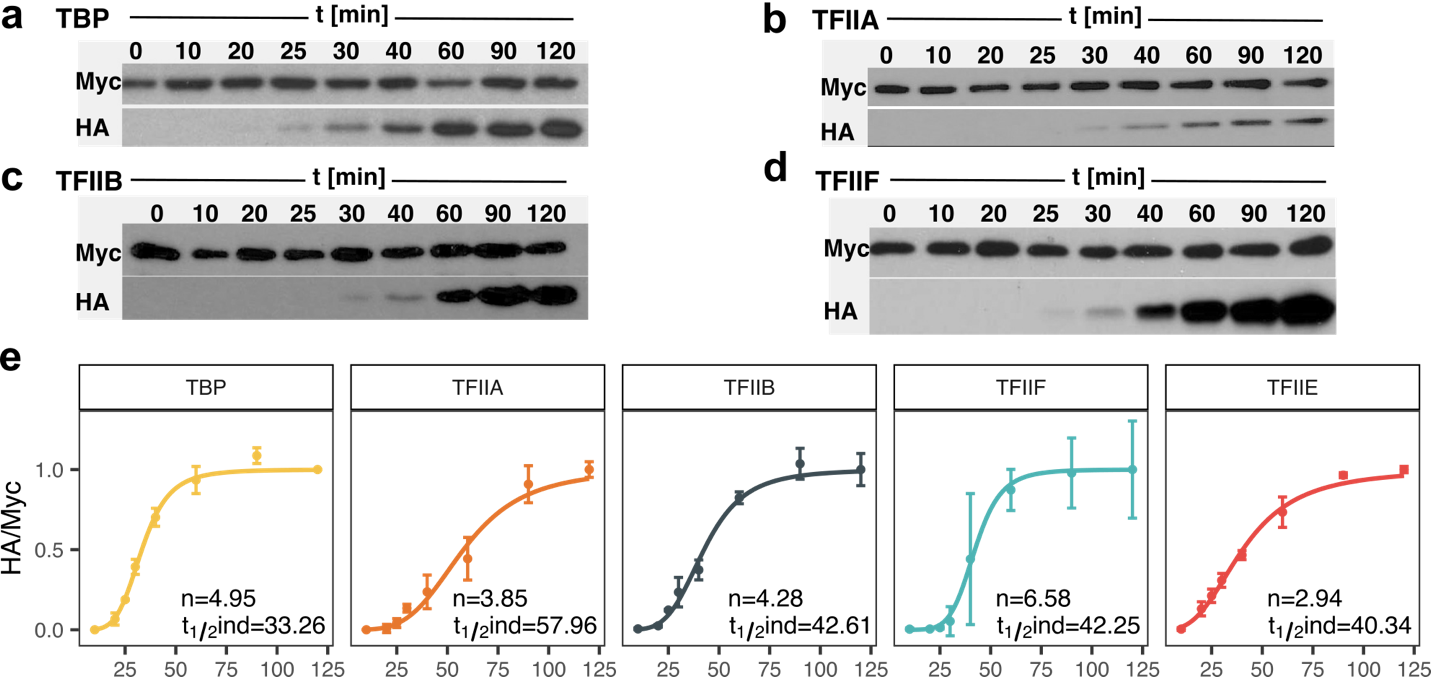

Fig. S1. Protein induction.

(a-d) Western blots of (a) TBP, (b) TFIIA (Toa1), (c) TFIIB, and (d) TFIIF (Tfg1) over the indicated time course. Myc tag indicates proteins made from genes expressed under control of their endogenous promoters, and the HA tag measures the level of the competitor expressed under galactose control. (e) Normalized HA/Myc ratios quantified from Western blots (n=3) with Hill fits. The error bars indicate standard deviations. Hill fit parameters are shown in the bottom right corner of each panel, n: Hill coefficients, t_1/2_ind: half time of HA-tagged protein induction.

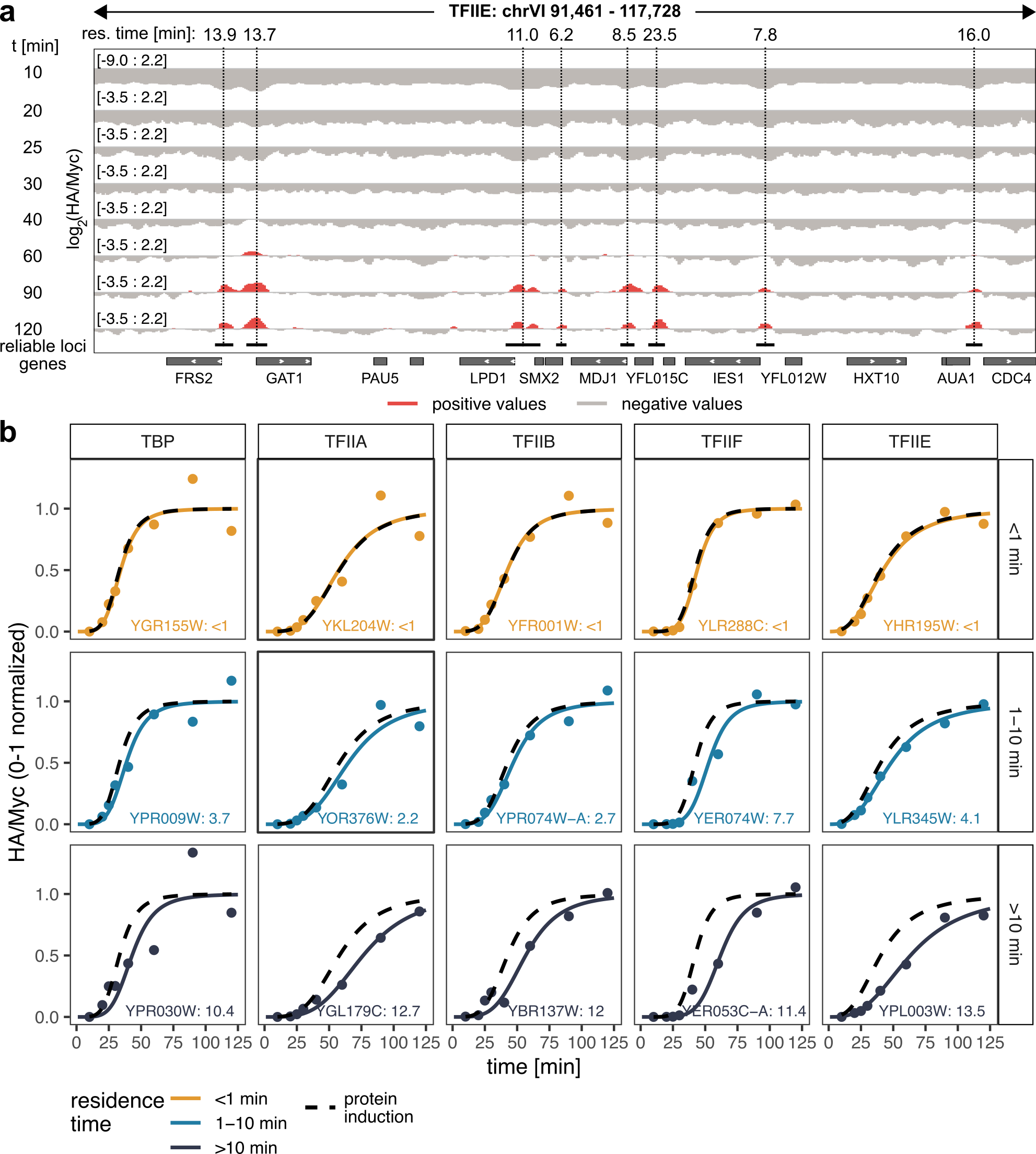

Fig. S2. Competition ChIP.

(a) Genome browser view showing log_2_ transformed ratios of normalized HA and Myc ChIP signals over time. (b) Examples of fitted ChIP-seq data for fast (<1 min), moderate (1-10 min) and slow sites (>10 min) for all GTFs examined in this study. Corresponding gene targets are indicated in the bottom right corner of each panel along with the estimated residence times in minutes. The plots show the normalized measurements of HA and Myc ChIP signal ratios fitted with colored curves; the dashed black curves show the induction rate for a given competitor, as also indicated in Fig. S1.

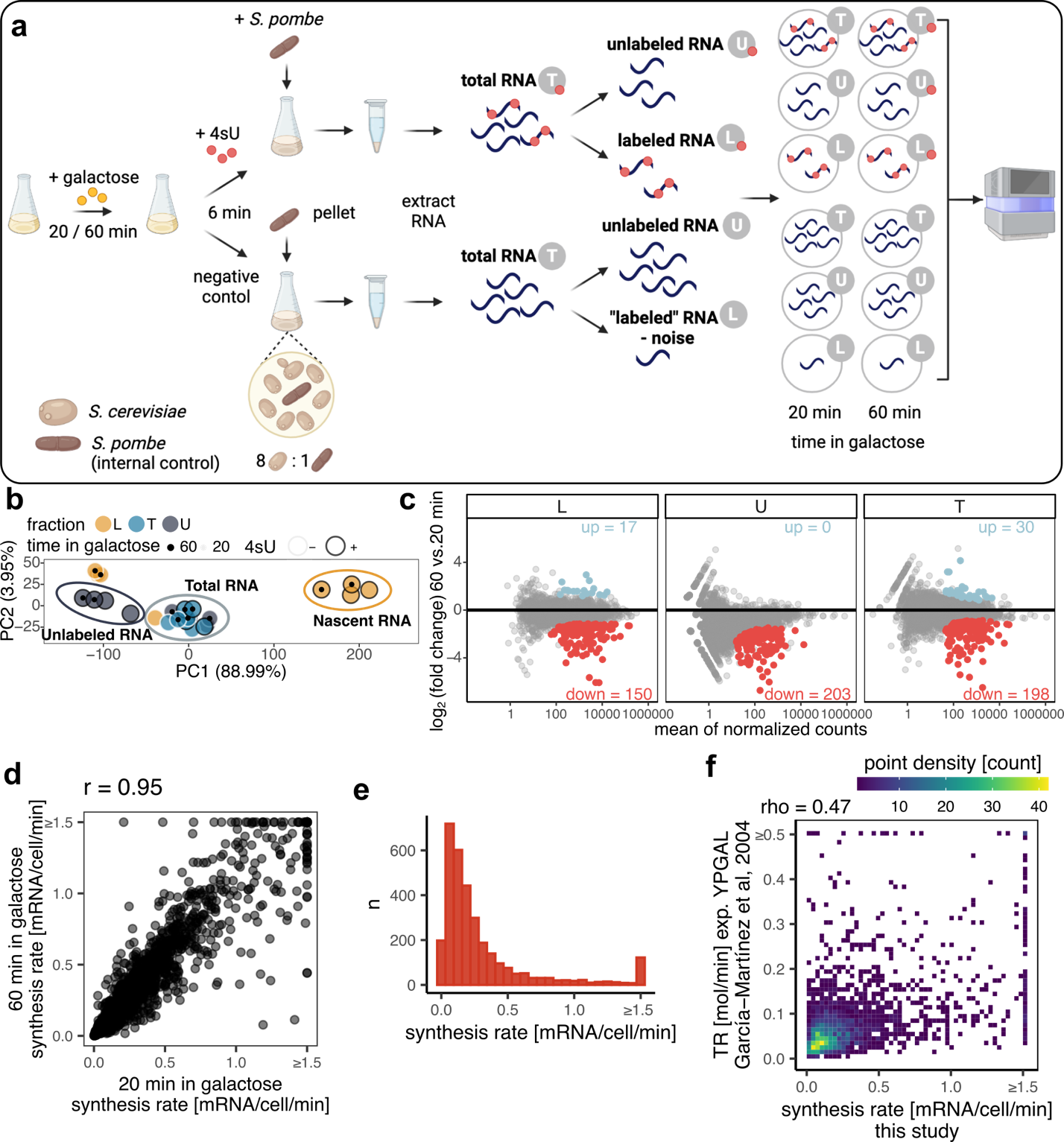

Fig. S3. Synthesis rate estimation with dynamic transcriptome analysis (DTA).

(a) Schematic overview of the DTA method as adapted in this study. (b) Principal component analysis (PCA) plot showing the first two principal components (PCs) calculated from normalized read coverage signal from all samples generated in this study. Highlighted are clusters of samples representing nascent RNA (L fraction after 4sU addition), total RNA (T fraction after 4sU addition as well as from negative control, along with U fraction from negative control), and unlabeled RNA (U fraction after 4sU addition). Percentages within the axis labels indicate percentage of variance explained by a given PC. (c) MA plot showing differentially expressed genes between samples grown for 60 vs. 20 minutes in galactose. Each point represents a gene, x-axis indicates size of a given gene in terms of the mean number of reads after normalization mapped to the gene, y-axis shows log_2_ of fold change between the two conditions. Highlighted are significantly misregulated genes, blue: upregulated at 60 minutes, red: downregulated at 60 minutes compared to the 20-minute time point, significance threshold: FDR-corrected p-value (padj) < 0.05. (d) Comparison of synthesis rates (in mRNA per cell per minute) estimated from samples grown for 20 minutes in galactose (x-axis) vs. 60 minutes in galactose (y-axis). Pearson’s correlation coefficient can be found above the plot. (e) Histogram showing the distribution of synthesis rates (in mRNA per cell per minute) estimated jointly from samples grown for 20 minutes and 60 minutes in galactose. Synthesis rates higher than 1.5 were combined into one bar to eliminate long tails. (f) Comparison of synthesis rates generated in this study (x-axis) to those generated by García-Martínez et al, 2004 (3). Spearman’s correlation coefficient is mentioned above the plot. Plot is color coded based on point density in each area. Symbol ≥ on the axis indicates that values higher than an indicated value were shrunk for plotting purposes to eliminate outliers.

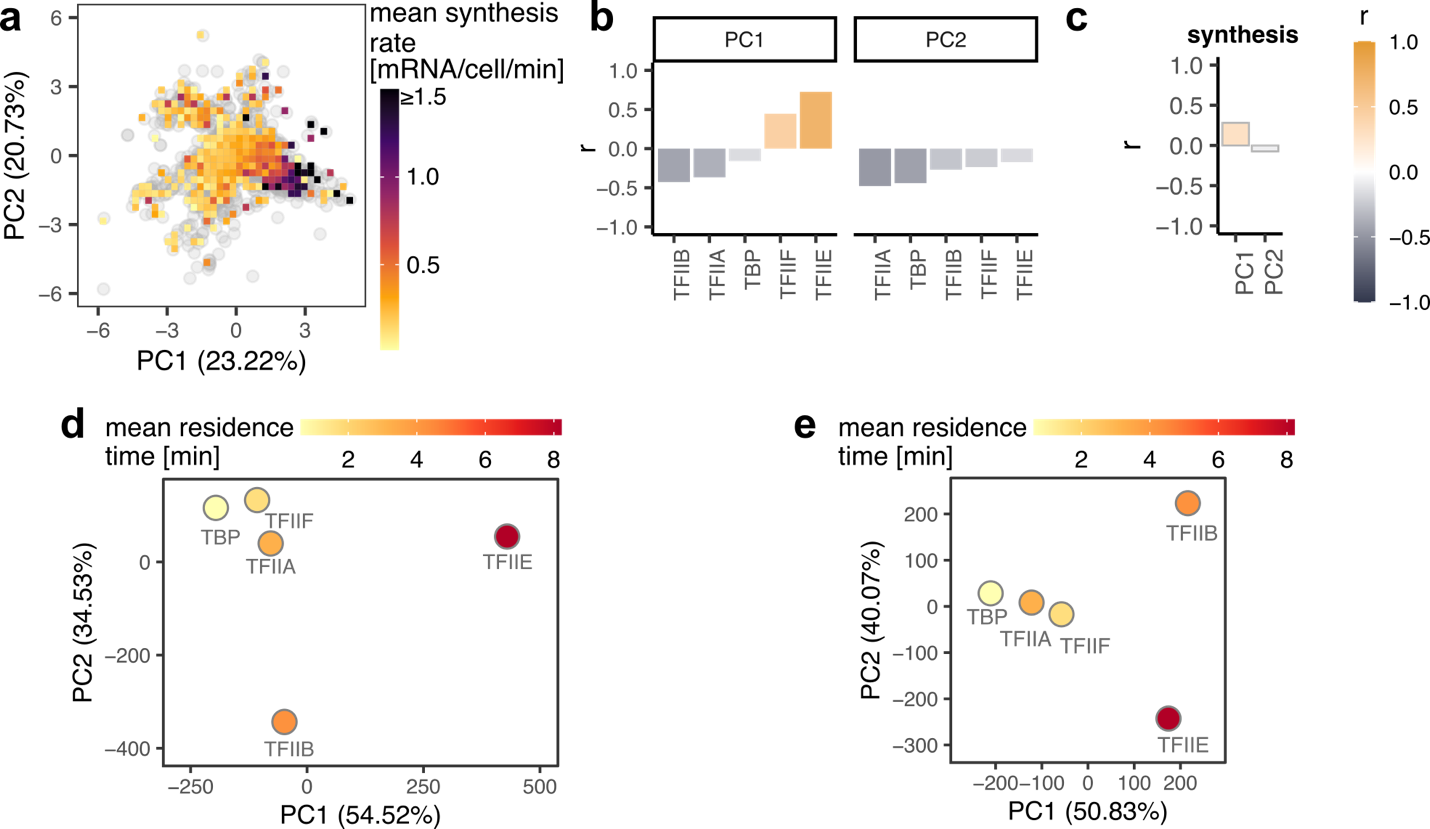

Fig. S4. Residence time PCA.

(a) PCA plot separating genes (points) based on GTF residence times. Color coding indicates the mean synthesis rate of the genes falling under a given area. Genes with residence time estimates <1 min were excluded from the analysis. (b) Person’s correlation coefficients (y-axis) between PCs from (a) and residence times of a given GTF (x-axis). (c) Pearson’s correlation coefficients (y-axis) between PCs (x-axis) from (a) and synthesis rates. (d) PCA plot separating individual GTFs (points) based on their residence times. (e) Similar to (d), however, sites with residence time estimates <1 min were excluded from the analysis. In PCA plots, the percentages in the axis labels indicate percentage of variance explained by a given PC.

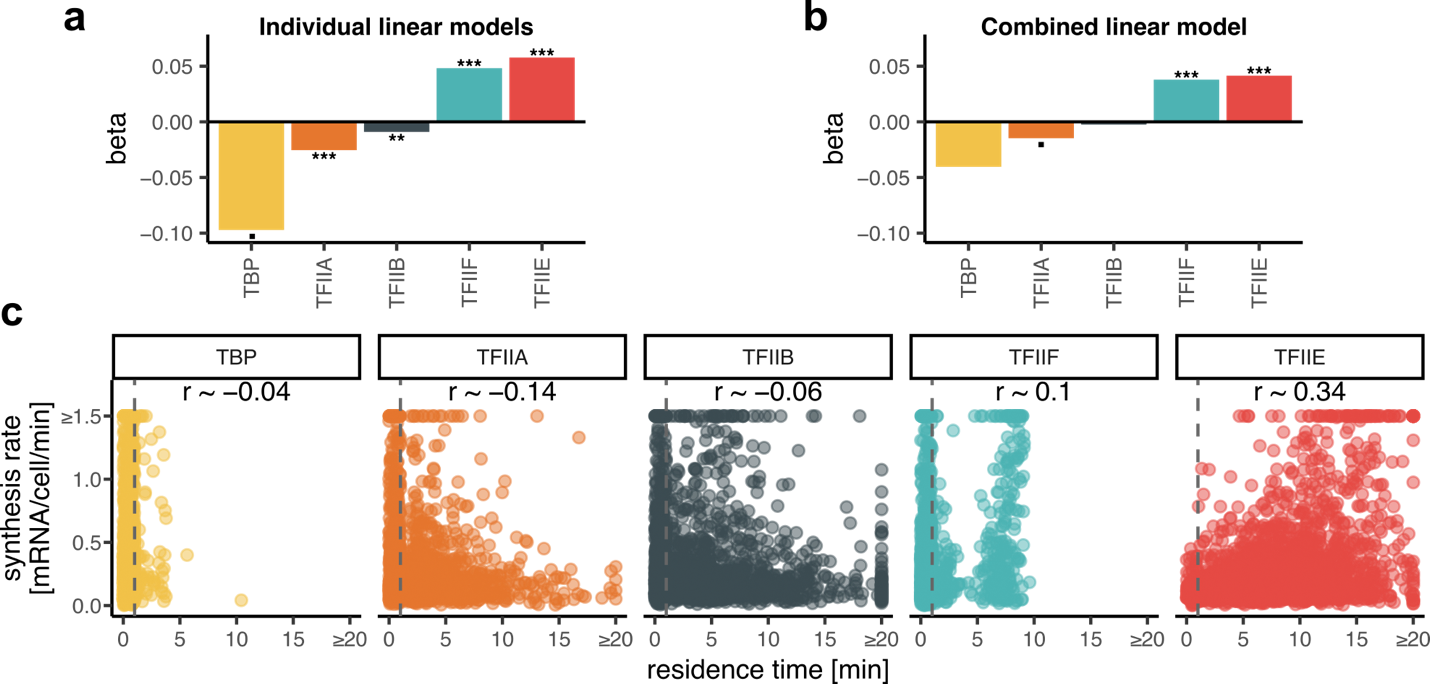

Fig. S5. Comparison between residence times and synthesis rates

(a) Bar plot showing β (beta) coefficients for linear models built between synthesis rates and residence times of an indicated GTF (synthesis rate ~ β*res. time_GTF_ + α). (b) Analogous to (b), showing coefficient from a linear model combining all factors (synthesis rate ~ β_TBP_*res. time_TBP_ + β_TFIIA_*res. Time_TFIIA_ + β_TFIIB_*res. Time_TFIIB_ + β_TFIIF_*res. Time_TFIIF_ + β_TFIIE_*res. Time_TFIIE_ + α). (c) Relationship between residence times (x-axis) and synthesis rates (y-axis) for GTFs as indicated. Pearson’s correlation coefficient estimates, r, are indicated in each panel. Symbol ≥ on the axis indicates that values higher than an indicated value were shrunk for plotting purposes to eliminate outliers. In the plots grey dashed line separates values randomly generated in this study for reliably fast sites. P-value symbols: . p < 0.1, * p < 0.05, ** p < 0.01, *** p < 0.001.

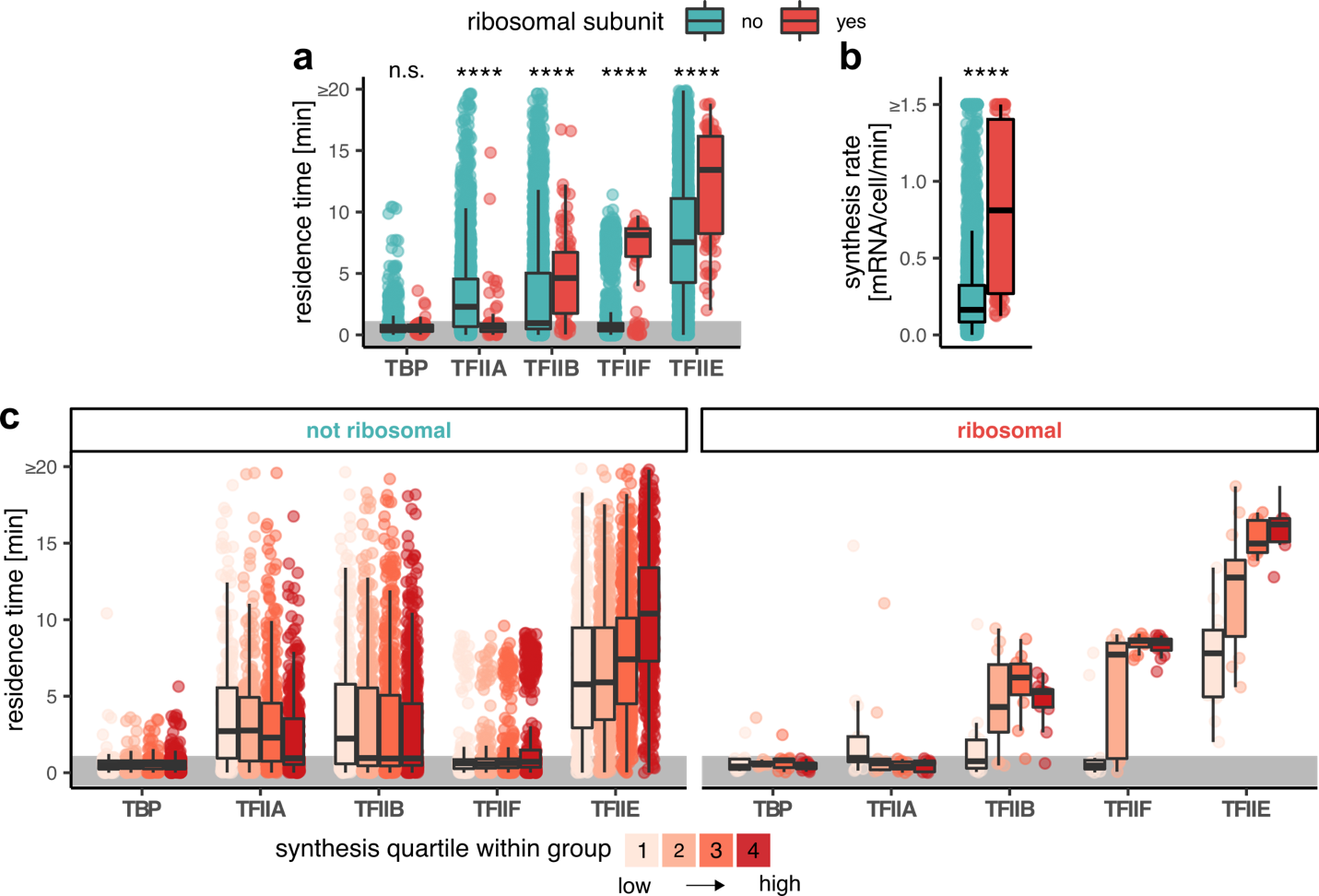

Fig. S6. Comparison of residence times between genes coding for ribosomal subunits and others.

(a) Box plots showing comparison of residence times (y-axis) for a given GTF (x-axis) for genes coding for ribosomal subunits (red) and other genes (green). (b) Box plot showing comparison of synthesis rates (y-axis) between genes coding for ribosomal subunits and other genes. (c) Box plots showing comparison of residence times (y-axis) for a given GTF (x-axis) across synthesis quartiles within genes coding for ribosomal subunits and other genes. In the plots grey area highlights values randomly generated in this study for reliably fast sites. Symbol ≥ on the axis indicates that values higher than an indicated value were shrunk for plotting purposes to eliminate outliers. P-value symbols: n.s. p > 0.05, * p <= 0.05, ** p <= 0.01, *** p <= 0.001, **** p <= 0.0001. In box plots the middle line represents the median, the lower and upper hinge represent the first and third quartiles, and the whiskers represent 1.5 * interquartile range.

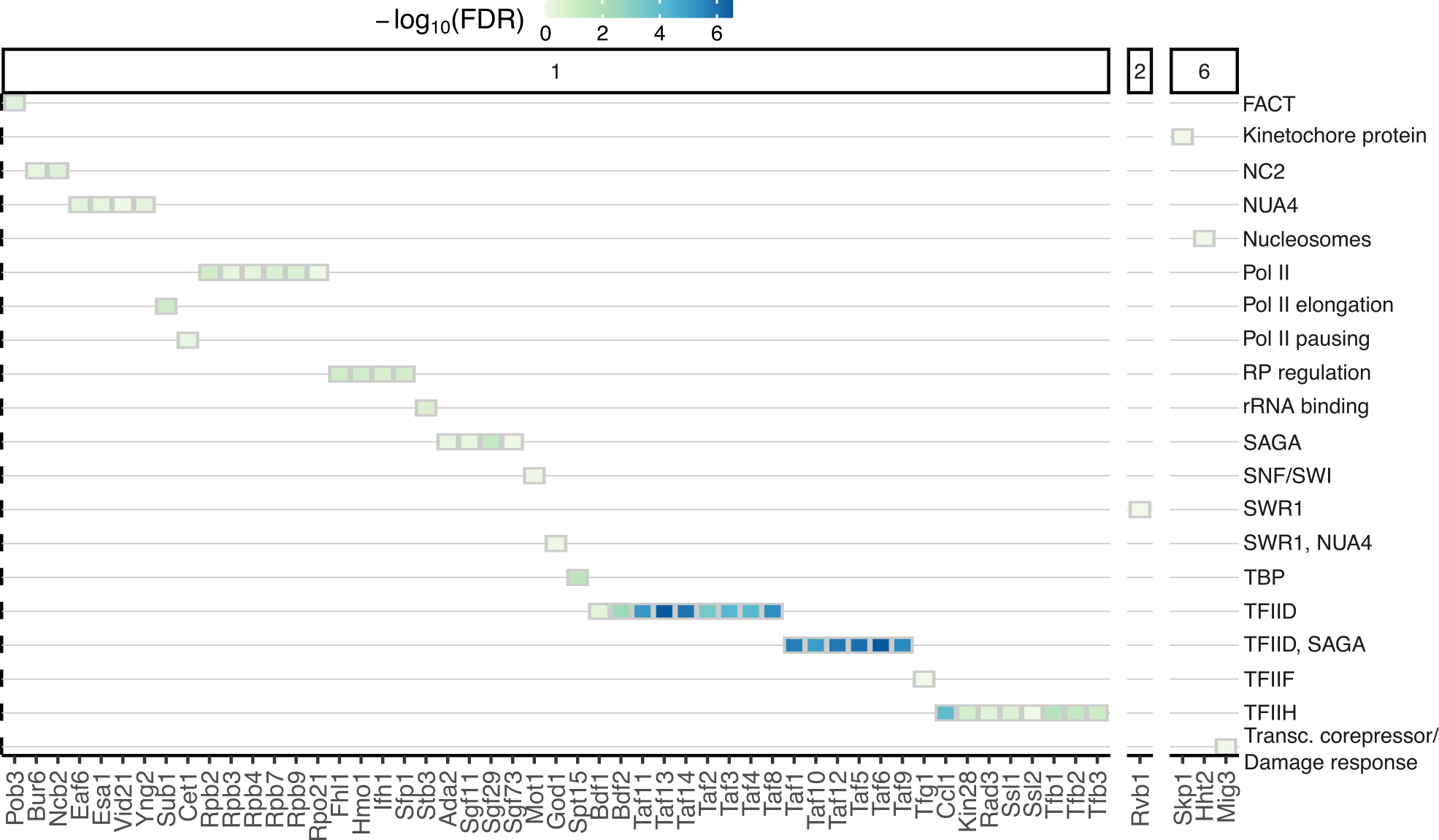

Fig. S7. Heatmap clusters enrichment.

The figure shows the yeast epigenome database transcription factors enriched in clusters in (a)/Fig. 5a classified into manually curated functional groups. Compared to Fig. 5d, subunits of GTFs and Pol II are included. The numbers on the top of the plot indicate the cluster number from (a)/Fig. 5a.

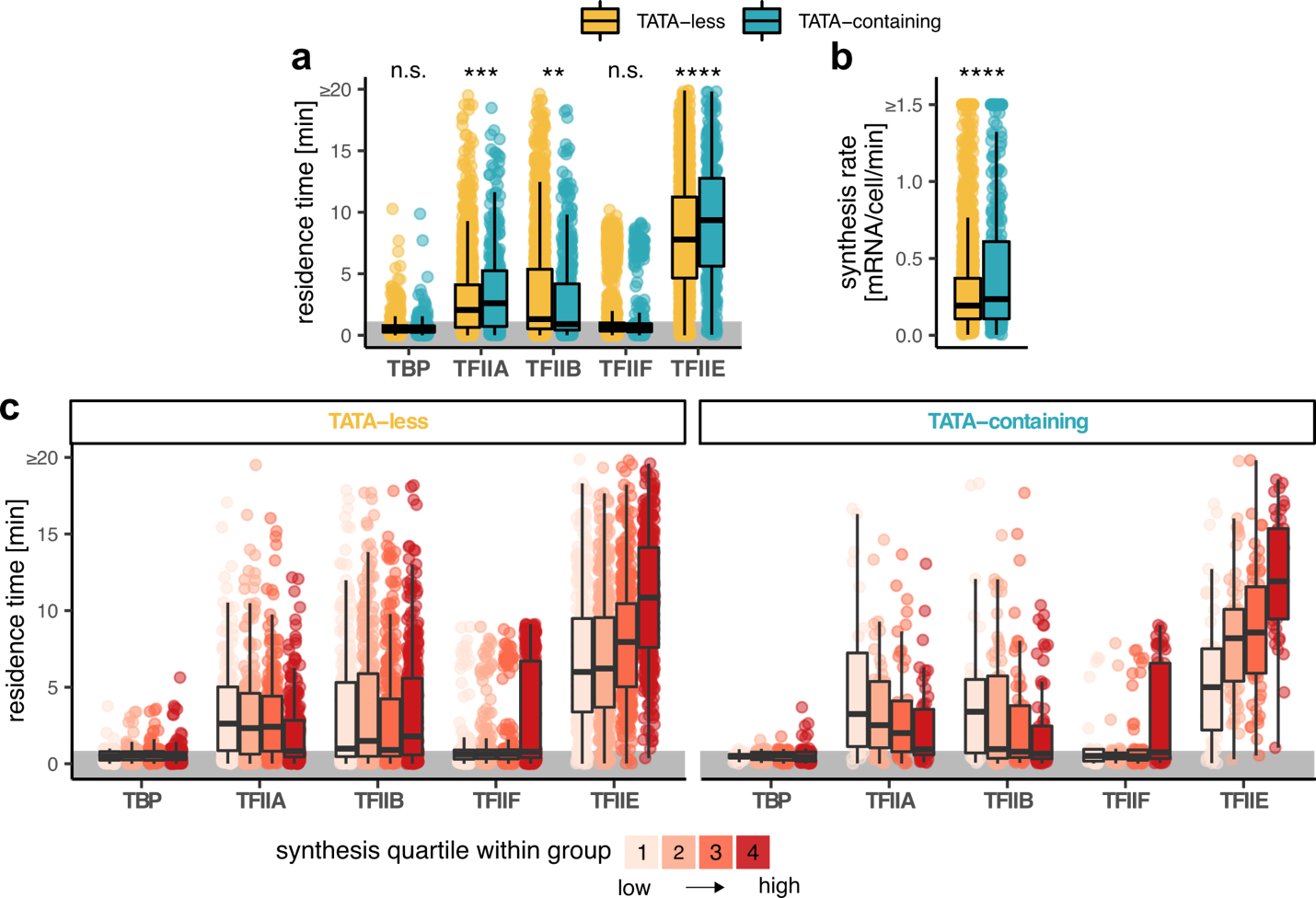

Fig. S8. Comparison of residence times between genes with or without a consensus TATA element in their promoters.

(a) Box plots showing comparison of residence times (y-axis) for a given GTF (x-axis) for genes without a consensus TATA element in their promoters (labeled TATA-less) and genes with a TATA-containing promoter (4). (b) Box plot showing comparison of synthesis rates (y-axis) between genes with TATA-less vs. TATA-containing promoters. (c) Box plots showing comparison of residence times (y-axis) for a given GTF (x-axis) across synthesis quartiles within genes with TATA-less (left panel) and TATA-containing (right panel) promoters. In the plots grey area highlights values randomly generated in this study for reliably fast sites. Symbol ≥ on the axis indicates that values higher than an indicated value were shrunk for plotting purposes to eliminate outliers. P-value symbols: n.s. p > 0.05, * p <= 0.05, ** p <= 0.01, *** p <= 0.001, **** p <= 0.0001. In box plots the middle line represents the median, the lower and upper hinge represent the first and third quartiles, and the whiskers represent 1.5 * interquartile range.

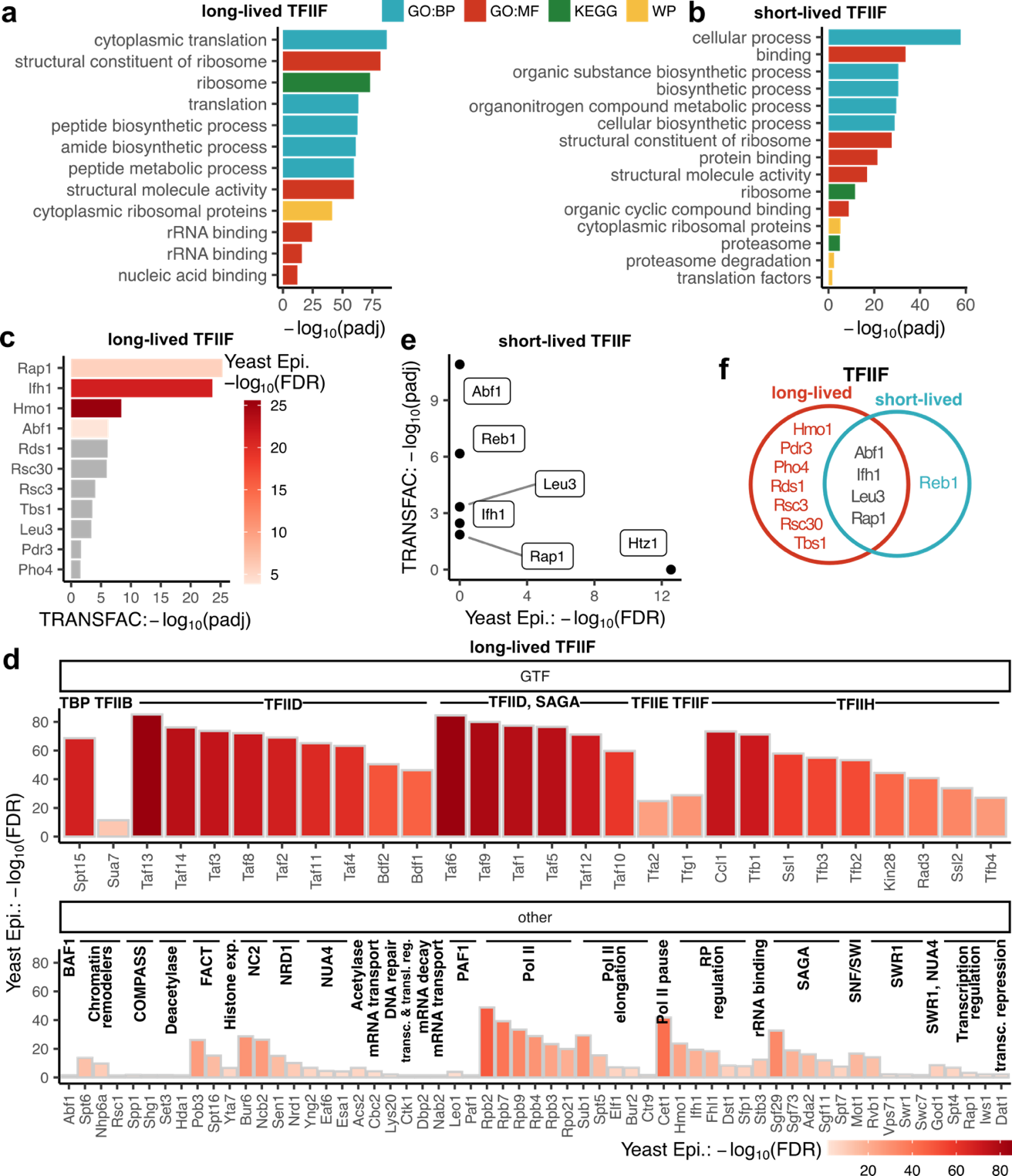

Fig. S9. Functional enrichment of long-lived and short-lived TFIIF sites.

As TFIIF long-lived sites were classified those with residence times ≥ 5 minutes, the rest were classified as short-lived. (a-b) Pathway enrichment of gene targets of long-lived (a) and short-lived (b) TFIIF sites. (c) TRANSFAC enrichment of long-lived TFIIF gene targets. TF enrichments also identified in Yeast Epigenome database are highlighted in shade of red based on Yeast Epigenome enrichment FDR-corrected p-values. (d) Enrichment of long-lived TFIIF gene targets in Yeast Epigenome DBF targets. Enriched factors are separated into general transcription factors (GTF) and others and were manually classified into categories. (e) Enrichment of short-lived TFIIF gene targets withing Yeast Epigenome database on x-axis vs enrichment within TRANSFAC. (f) Venn diagram showing TRANSFAC TFs enriched in long-lived vs. short-lived TFIIF sites. GO:BP GO biological process, GO:MF GO molecular function, WP WikiPatways. Significant enrichment: FDR p < 0.05. See Methods section “Yeast DBF database (Yeast Epigenome)” for more details on Yeast Epigenome database.

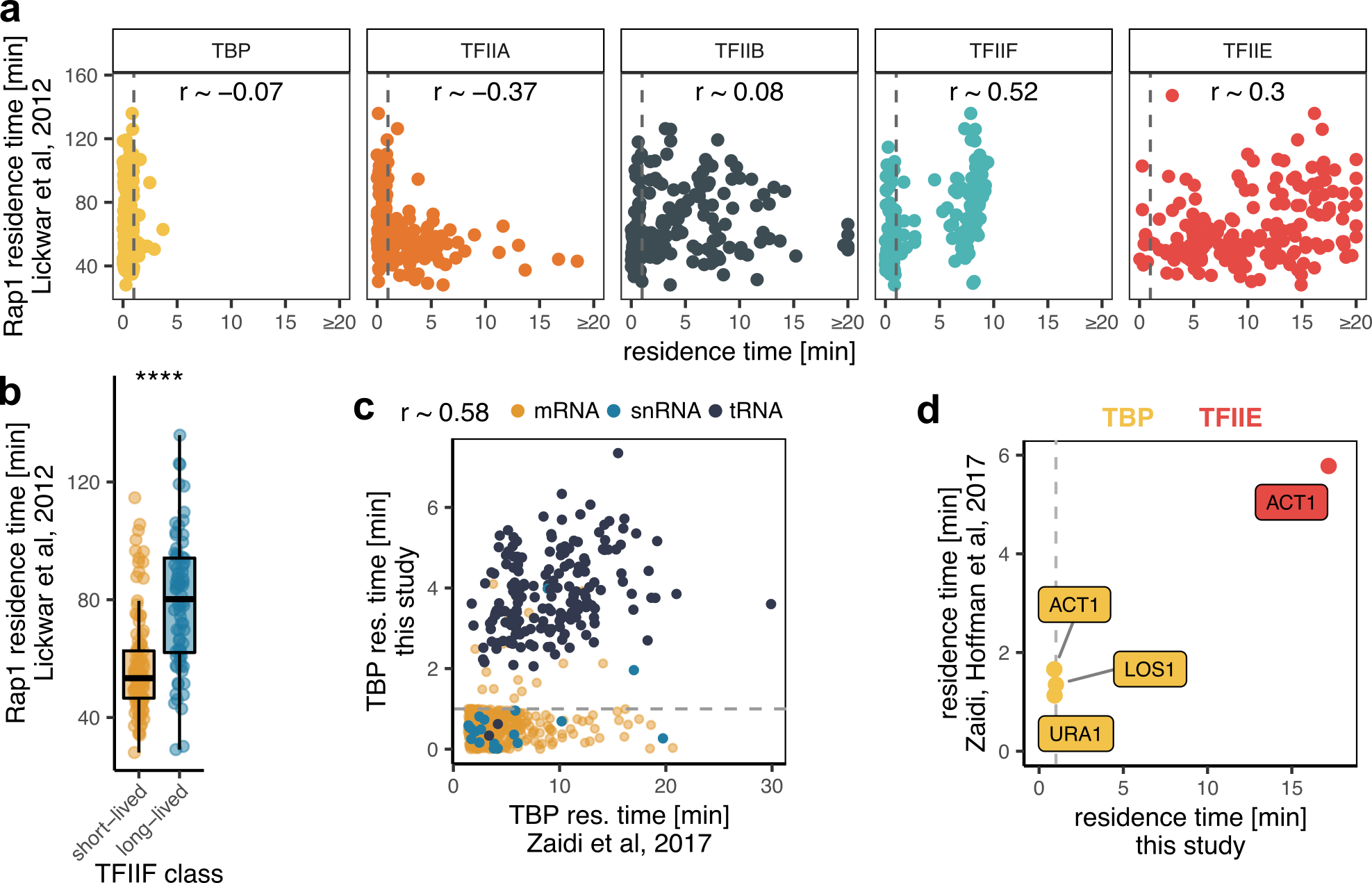

Fig. S10. Comparison to Rap1 residence times and comparison of TBP and TFIIE residence time s to others.

(a) Comparison between residence times of GTFs from this study (x-axis) to Rap1 residence times from Lickwar et al, 2012 (5). Pearson’s correlation coefficient estimates, r, are indicated in each panel. Grey dashed line separates randomly generated residence time values for reliably fact sites (<1 min). (b) Box plots showing the difference in Rap1 residence times at gene targets of short-lived (<=5 minutes) vs. long-lived (>5 minutes) TFIIF sites. (c) Comparison of TBP residence time estimates from this study (y-axis) to those from Zaidi et al, 2017 (1) (x axis). Pearson’s correlation coefficient estimate is indicated above the plot. (d) Comparison of TBP and TFIIE residence time estimates from this study (x-axis) to those from Zaidi, Hoffman, et al, 2017 (2) (y-axis). In the plots grey dashed line separates values randomly generated in this study for reliably fast sites. Symbol ≥ on the axis indicates that values higher than an indicated value were shrunk for plotting purposes to eliminate outliers. In the box plots the middle line represents the median, the lower and upper hinge represent the first and third quartiles, and the whiskers represent 1.5 * interquartile range.

### Supplementary references

1.	Zaidi HA, Auble DT, Bekiranov S. RNA synthesis is associated with multiple TBP-chromatin binding events. Sci Reports 2017 71. 2017 Jan 4;7(1):1–12.

2.	Zaidi H, Hoffman EA, Shetty SJ, Bekiranov S, Auble DT. Second-generation method for analysis of chromatin binding with formaldehyde–cross-linking kinetics. J Biol Chem. 2017 Nov 24;292(47):19338–55.

3.	García-Martínez J, Aranda A, Pérez-Ortín JE. Genomic Run-On Evaluates Transcription Rates for All Yeast Genes and Identifies Gene Regulatory Mechanisms. Mol Cell. 2004 Jul 23;15(2):303–13.

4.	Basehoar AD, Zanton SJ, Pugh BF. Identification and Distinct Regulation of Yeast TATA Box-Containing Genes. Cell. 2004 Mar 5;116(5):699–709.

5.	Lickwar CR, Mueller F, Hanlon SE, McNally JG, Lieb JD. Genome-wide protein–DNA binding dynamics suggest a molecular clutch for transcription factor function. Nat 2012 4847393. 2012 Apr 11;484(7393):251–5.
